# Cohesin depleted cells rebuild functional nuclear compartments after endomitosis

**DOI:** 10.1101/816611

**Authors:** Marion Cremer, Katharina Brandstetter, Andreas Maiser, Suhas S P Rao, Volker Schmid, Miguel Guirao-Ortiz, Namita Mitra, Stefania Mamberti, Kyle N Klein, David M Gilbert, Heinrich Leonhardt, Maria Cristina Cardoso, Erez Lieberman Aiden, Hartmann Harz, Thomas Cremer

**Affiliations:** Anthropology and Human Genomics, Department Biology II, Ludwig-Maximilians-Universität München, Germany; Human Biology & BioImaging, Center for Molecular Biosystems, Department Biology II, Ludwig-Maximilians-Universität München, Germany; Center for Genome Architecture, Department of Molecular and Human Genetics, Baylor College of Medicine, Houston, Texas, United States of America; Department of Structural Biology, Stanford University School of Medicine, California, United States of America; BioImaging Group, Department of Statistics, Ludwig-Maximilians-Universität München, Germany; Cell Biology and Epigenetics, Department of Biology, Technische Universität Darmstadt, Germany; Department of Biological Science, Florida State University, Tallahassee, Florida, United States of America; Center for Theoretical Biological Physics, Rice University, Houston, Texas, United States of America; Broad Institute of the Massachusetts Institute of Technology and Harvard University, Cambridge, Massachusetts, United States of America; Departments of Computer Science and Computational and Applied Mathematics, Rice University, Houston, Texas, United States of America

## Abstract

Cohesin plays an essential role in chromatin loop extrusion, but its impact on a compartmentalized nuclear architecture, linked to nuclear functions, is debatable. Using live-cell and super-resolved 3D microscopy, we demonstrate that cohesin depleted cells pass through an endomitosis and rebuild a single multilobulated nucleus (MLN) with chromosome territories (CTs) pervaded by interchromatin channels. CTs contain chromatin domain clusters with a zonal organization of repressed chromatin domains in the interior and transcriptionally competent domains located at the periphery. Splicing speckles are located nearby within the lining channel system. These clusters form microscopically defined, active and inactive compartments, which correspond to A/B compartments, detected with ensemble Hi-C. Functionality of MLN despite continuous absence of cohesin was demonstrated by their ability to pass through S-phase with typical spatio-temporal patterns of replication domains. Evidence for structural changes of these domains compared to controls suggests that cohesin is required for their full integrity.

## Introduction

Cohesin, a ring-like protein complex with its major subunits RAD21, SMC1 and SMC3 exerts its key functions by tethering distant genomic loci into chromatin loops. It is involved in sister chromatid entrapment to ensure proper chromosome segregation during mitosis, in double strand break repair and gene regulation, and importantly was found essential for chromatin loop extrusion by shaping loops in the sub-Mb range anchored at CTCF/cohesin binding sites ^1-4, 5,6^ for review see ^7-13^.

These results have argued for an essential role of cohesin in the formation of a functional nuclear architecture. Studies of the impact of cohesin depletion on nuclear structure and function have become greatly facilitated by an auxin-inducible degron (AID) system, which triggers a rapid and selective proteolysis of RAD21 after addition of auxin to the culture medium resulting in the loss of cohesin from chromatin ^14^. Using this system in the colon cancer derived HCT116-RAD21-mAC cell line, we previously demonstrated the rapid disappearance of chromatin loop domains with a concomitant loss of topologically associated domains (TADs) in Hi-C contact matrices averaged over large cell populations, with only minor effects of cohesin depletion on gene expression ^15^. Other studies, using different cell types and approaches for cohesin elimination yielded similar results, reviewed in ^16^.

Here, we studied the long-term fate of cohesin depleted cells at the single cell level with live-cell and super-resolved quantitative microscopy and performed a thorough comparison with Hi-C and related Repli-seq ^17,18^ methods. These approaches complement each other in ways that cannot be achieved by either method alone. Unexpectedly, we observed that cohesin depleted interphase cells are able to pass through an endomitosis yielding a single postmitotic cell with a multilobulated cell nucleus (MLN). MLN formation was accompanied by the rebuilding of chromosome territories (CTs) and the reconstitution of functional A and B compartments detected by ensemble Hi-C experiments, as well as co-aligned active and inactive nuclear compartments (ANC / INC) based on microscopic studies, reviewed in ^19,20^. In line with these principal features of a functional nuclear architecture, we found in our present study that MLN are able to initiate and traverse through S-phase with typical stage specific patterns of replication domains (RDs). Quantitative 3D image analyses indicated a larger number of RDs together with an increased heterogeneity of RD volumes. TADs, however, remained missing in ensemble Hi-C studies of cohesin depleted MLN. Our findings demonstrate the maintenance of spatial arrangements of RDs in the absence of cohesin and also support a role of cohesin in the compaction of functional higher order chromatin structures ^21^. A joint presentation of results from quantitative 3D microscopy and Hi-C studies is complicated by a different terminology to describe the structural and functional higher order chromatin entities discovered by either approach. For a definition of terms as we use them below, we refer readers to Supplementary Table 1.

## Results

### Validation of auxin induced proteolysis of the cohesin subunit RAD21

All experiments of this study were performed with the human colon cancer derived cell line HCT116-RAD21-mAC ^14^, where an auxin-inducible degron (AID) is fused to both endogenous RAD21 alleles together with a sequence coding for a fluorescent reporter (see Supplementary Fig.1). About 98% of nuclei in untreated control cell cultures expressed RAD21-mClover. Selective degradation of RAD21 under auxin treatment (6h in 500 μM auxin) was shown by negative immunostaining with a RAD21 antibody, while epitopes of cohesin subunits SMC1 and SMC3 remained intact under auxin (Supplementary Fig. 2A). RAD21-mClover degradation was quantitatively assessed by intensity measurements recorded from high throughput imaging of single cells after 6h auxin treatment (Supplementary Fig. 2B). A visible decline of RAD21-mClover fluorescence was first noted in time lapse images 30 min after incubation of cells in 500 μM auxin and appeared completed within 4:00h (Supplementary Fig. 3A). Furthermore, quantitative measurements of RAD21-mClover decline over time were performed on a single cell level (for details see Supplementary Fig. 3B-C). Notably, ∼4% of cells escaped auxin induced RAD21 degradation. In order to exclude non-responsive cells from further analyses of the impact of cohesin depletion, RAD21-mClover fluorescence was routinely recorded in all experiments with auxin treated cell populations except for 3D-FISH experiments where DNA heat denaturation degrades the reporter fluorescence ^22^.

**Fig. 1:**
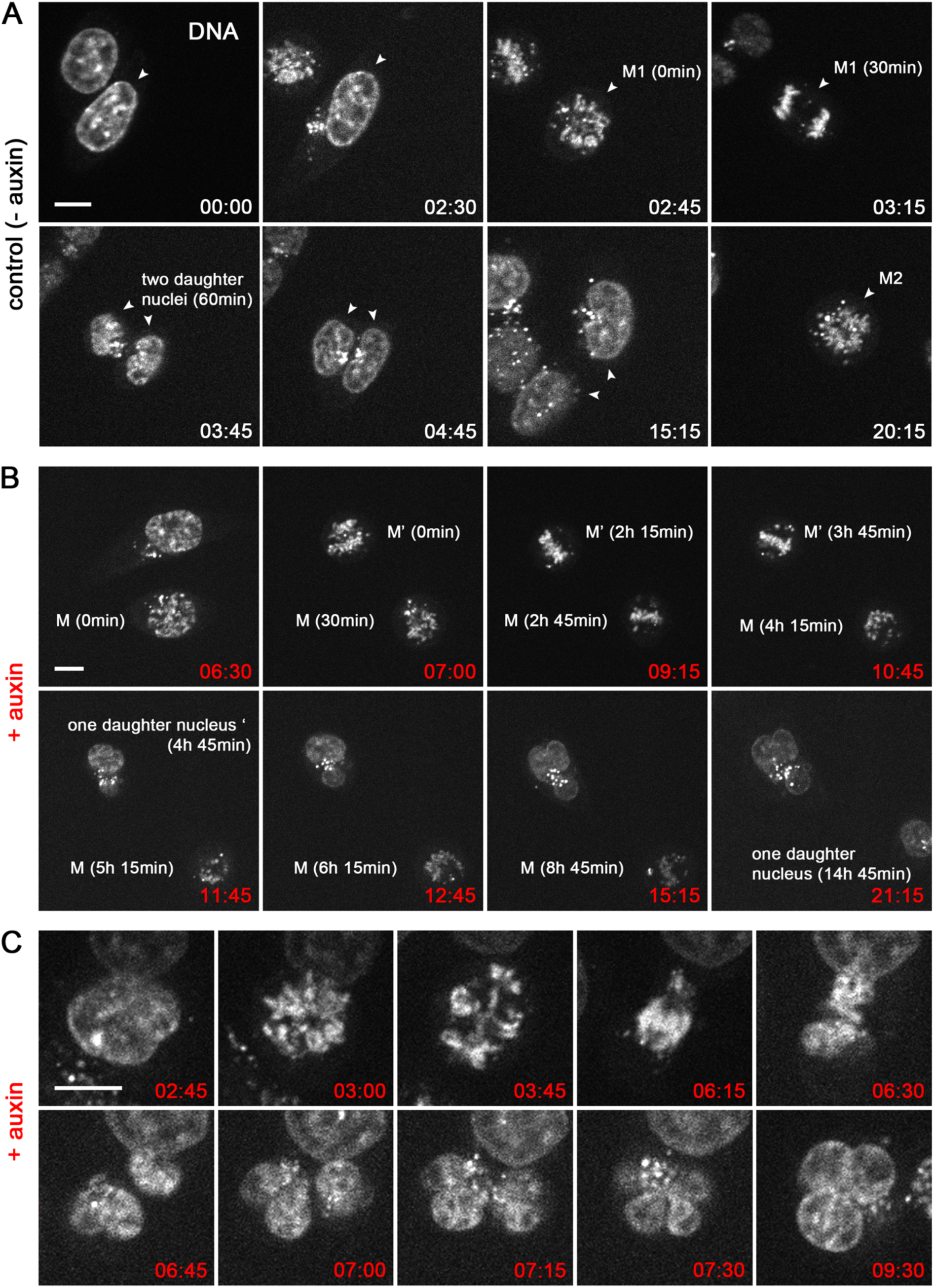
Live cell microscopy demonstrating prolonged abnormal mitosis with subsequent formation of one endomitotic multilobulated nucleus (MLN) in cohesin depleted cells. **(A)** Selected points from time lapse imaging (Σt=21h, Δt=15min) of untreated control cells (DNA stained with SiR-DNA) with accomplishment of mitosis (M1) within 1h (time 02:45 – 03:45) and subsequent formation of two daughter nuclei. A second mitosis (M2) of one daughter nucleus is shown at time 20:15. **(B)** Selected time lapse images of nuclei after cohesin degradation conducted in parallel to control cells demonstrate a prolonged mitotic stage. Mitosis (M) emerges at time 6:30 after auxin treatment, transition into one abnormal multilobulated daughter nucleus (MLN) is seen 14:45h later (time 21:15). Mitosis (M’) emerges 7h after auxin treatment (time 07:00), transition into an MLN is seen 4:45h later (time 11:45). **(C)** Time lapse imaging from the same series at a higher zoom shows an aberrant mitosis with an adumbrated formation of two daughter nuclei (time 06:45), that finally appear as one MLN at time 7:15. Scale bar: 10 µm. The complete series of time lapse images shown in A-C and raw data of additional observations are provided in https://cloud.bio.lmu.de/index.php/s/rZxxkgYExonWLgy?path=%2FFig1

**Fig. 2:**
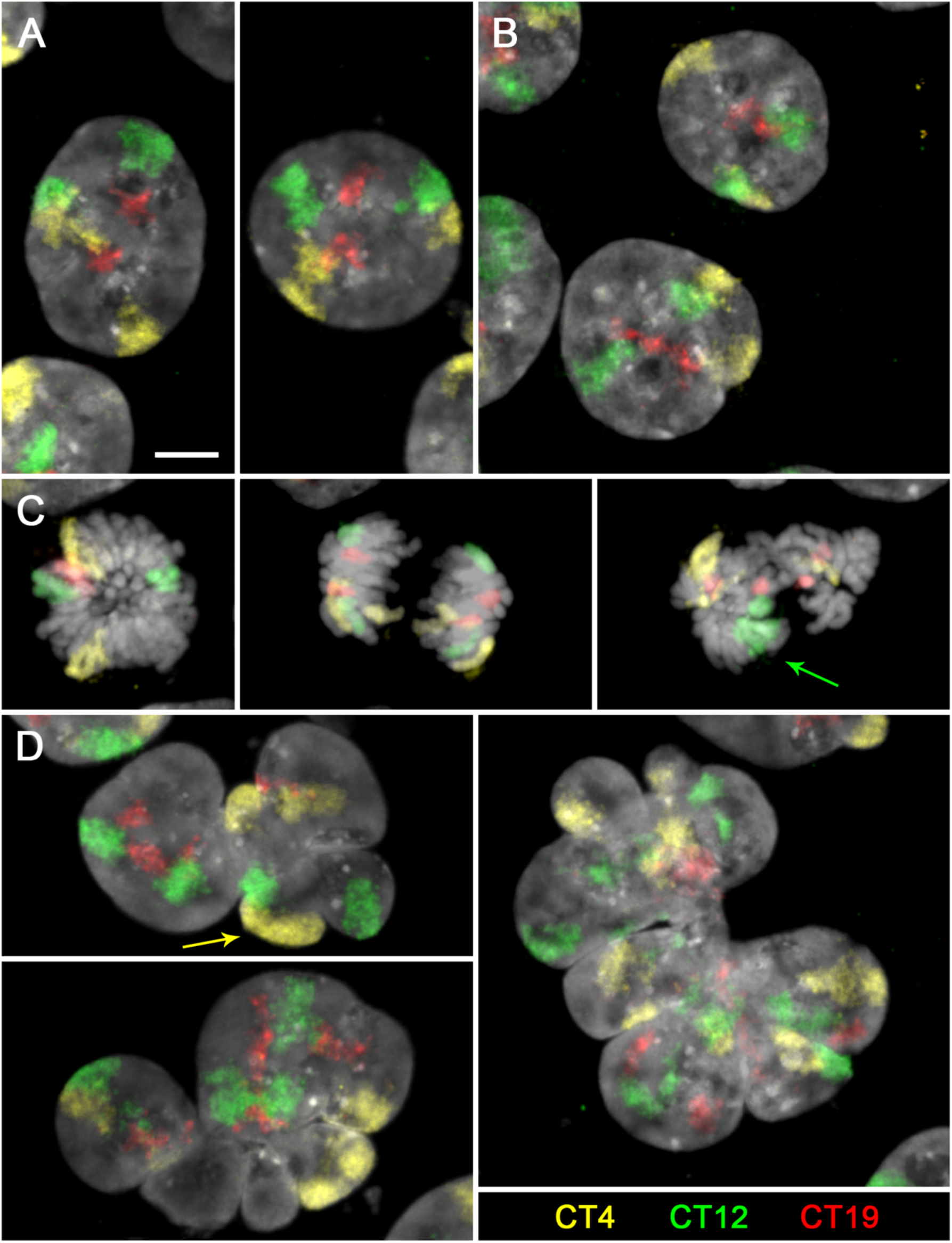
Maintenance of chromosome territories (CTs) in cohesin depleted nuclei and their reconstitution after endomitosis. **(A-E)** Z-projections of entire DAPI stained nuclei (gray) with painted territories of chromosomes 4 (yellow), 12 (green) and 19 (red) acquired by confocal fluorescence microscopy. **(A)** control nuclei and **(B)** pre-mitotic cohesin depleted nuclei after 6h in auxin show two inconspicuous copies for each CT. **(C)** Mitoses from 6h auxin treated cultures with two coherent chromosomes in a (presumably early) metaphase plate *(left)*, after chromatid segregation (mid) and missegregation of chromosome 12 (arrow) in an abnormal mitotic figure *(right)*. **(D)** *left:* two endomitotic multilobulated nuclei (MLN) with four copies for each CT. Arrow marks two CTs 4 that are overlayed in the z-projection. *Right:* Large MLN with a torn-up appearance of CTs with seemingly >4 painted regions for each CT (compare also Supplementary Fig. 5). Scale bar: 5 µm. Z-stacks of nuclei shown in A-D and z-projections of nuclei from additional experiments are provided in https://cloud.bio.lmu.de/index.php/s/rZxxkgYExonWLgy?path=%2FFig2

**Fig. 3:**
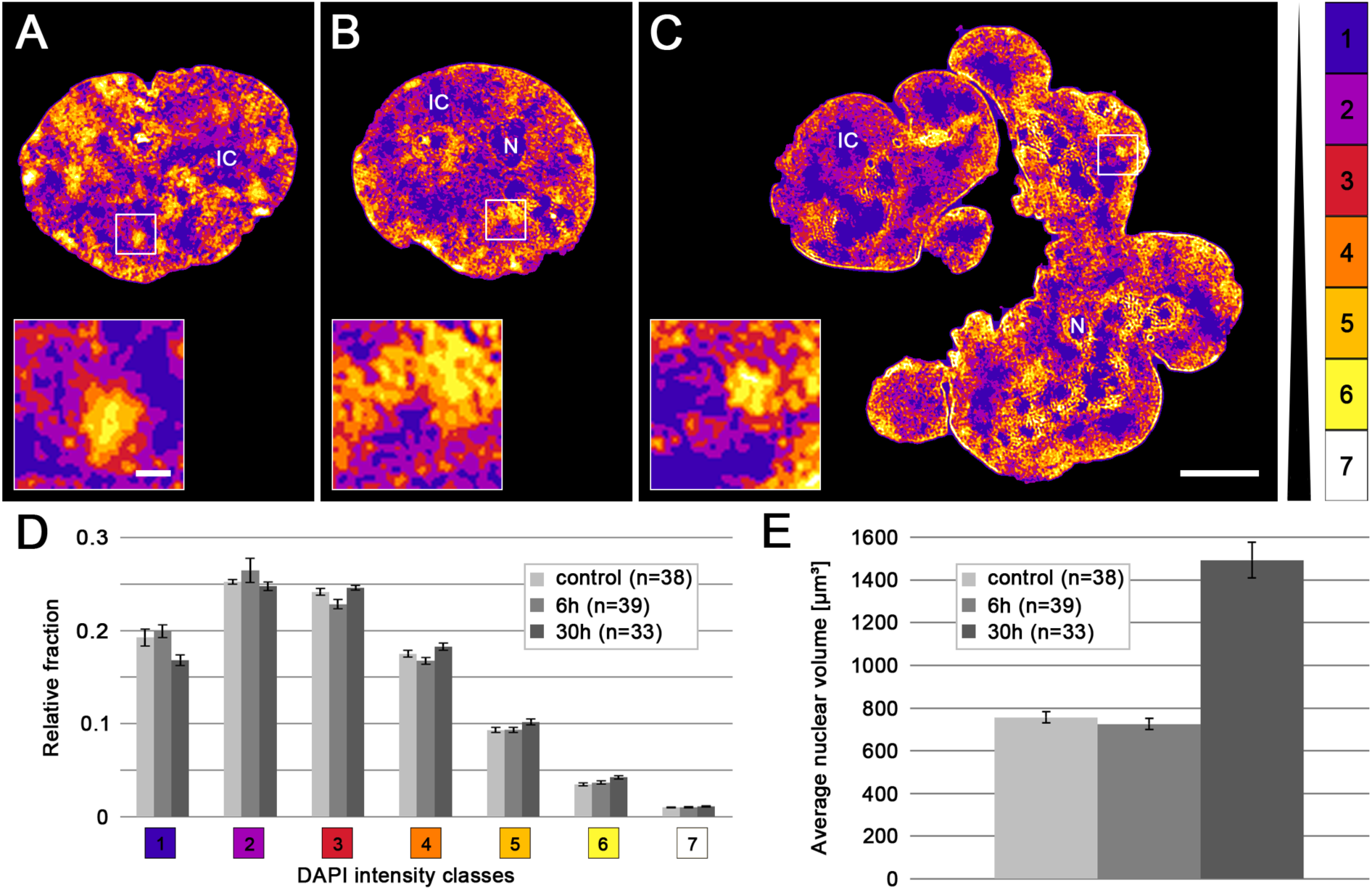
Compartmentalized architecture with an IC channel network pervading chromatin domain clusters (CDC) with zonal compaction differences in controls and cohesin depleted, pre- and post-endomitotic nuclei. **(A-C)** DAPI stained mid-sections of representative nuclei acquired by 3D-SIM from **(A)** control nucleus; **(B)** cohesin depleted nucleus (6h auxin); **(C)** cohesin depleted multilobulated nucleus (MLN) (30h auxin) are displayed by seven DAPI intensity classes in false colors, used as proxies for chromatin compaction ^29^. Class 1 (*blue*) pixels close to background intensity, largely reflecting the interchromatin compartment (IC) with only sparse DNA, class 7 (*white*) pixels with highest intensities (color code on the right). All nuclei in A-C reveal a network of chromatin domain clusters (CDCs) comprising a compacted core and a surrounding low-density zone co-aligned with class 1 regions that meander between CDCs as part of the IC (see insets). Likewise, all nuclei display a rim of compacted (hetero)chromatin at the nuclear periphery. N = nucleolus; IC = interchromatin channels/lacunae. Scale bars: 5 µm, insets: 0.5 µm **(D)** Relative 3D signal distributions of DAPI intensity classes in control nuclei and cohesin depleted nuclei show an overall similar profile. **(E)** Average nuclear volumes from the same series of nuclei. The ∼2-fold increase of nuclear volumes in (post-endomitotic) MLN after 30h auxin likely reflects their further increase of a 2n DNA content immediately after endomitosis to a 4n DNA content after another round of DNA replication (compare Fig. 6 E), for statistical tests see Supplementary Table 3. Complete image stacks from nuclei shown in A-C, data for DAPI intensity classifications and nuclear volumes in individual nuclei are provided in https://cloud.bio.lmu.de/index.php/s/rZxxkgYExonWLgy?path=%2FFig3

### Cohesin depleted cells pass through a prolonged endomitosis yielding a daughter cell with one multilobulated nucleus (MLN)

Using time lapse imaging over 21h at Δt=15min, we compared entrance into mitosis, mitotic progression and exit in parallel in untreated controls and in cohesin depleted cells, where auxin was added just before starting live cell observations. In control cells ∼80% of all recorded mitoses (n=45) passed mitosis within <1h and formed two inconspicuous daughter nuclei. A second mitosis observed for individual nuclei ∼20h after the first division demonstrates their capacity to divide again under the given observation conditions (Fig. 1A). Notably, about 20% of mitoses recorded in untreated control cells revealed prolonged mitoses (>2h) followed by transition into an abnormal cell nucleus (for detailed information on individual nuclei see Supplementary Table 2), a feature which is not unusual in tumor cell lines (reviewed in ^23^). In cohesin depleted cells (n=36) mitotic entrance was inconspicuous (Fig.1B), mitotic progress, however, was consistently delayed up to 14h (median 4.5h, for detailed information on individual nuclei see Supplementary Table 2). This prolonged mitotic stage raised the mitotic index in cohesin depleted cell cultures after 6h in auxin to almost 30% versus ∼4% in control cultures (Supplementary Fig. 4). The delayed mitotic passage was associated with the formation of abnormal, e.g. multipolar mitotic figures persisting over several hours (Fig.1B). Fig. 1C depicts a mitotic cell apparently approaching the stage of two separated daughter nuclei. Despite their seemingly complete separation, these daughter nuclei were presumably still connected by filaments (see below and Supplementary Fig. 5) and did not complete karyokinesis. All cohesin depleted cells that were followed through an entire mitosis (n=23, Supplementary Table 2) resulted in the formation of a single multilobulated nucleus (MLN) within one daughter cell, indicative for an endomitotic event ^24^. As a consequence, in cell cultures fixed 30h after cohesin depletion, MLN accumulated up to ∼60% versus ∼2% in control cultures (Supplementary Fig.4). After 50h in cell culture, the number of MLN declined due to increased apoptosis (data not shown).

**Fig. 4:**
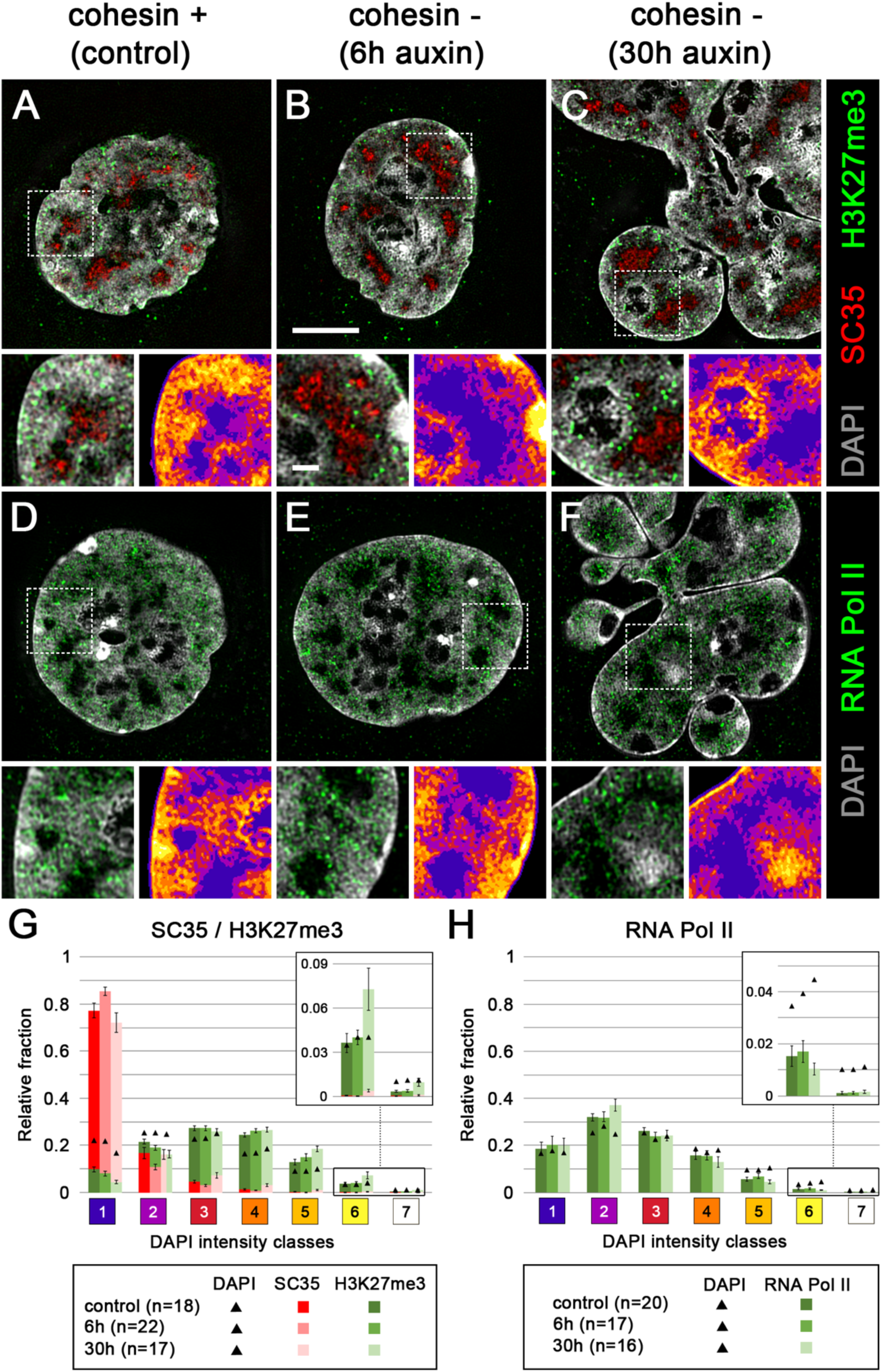
Nuclei of control and cohesin depleted cells show a congruent 3D topography of SC35, H3K27me3 and RNA Pol II. **(A-F**) DAPI stained nuclear mid-sections (gray) displayed from 3D SIM image stacks of control nuclei (A,D), pre-mitotic, cohesin depleted nuclei after 6h auxin treatment (B,E), and post-endomitotic MLN after 30h auxin treatment reveal the topography of immunostained SC35 (red) and H3K27me3 (green) (A-C), or active RNA Pol II (red) (D-F). Scale bar: 5 µm. An enlargement of a representative, boxed area is shown (left) beneath each mid-section (left side), together the color-coded DAPI intensity heat map (right) (compare Fig. 3). Scale bar: 1 µm. SC-35 marked splicing speckles are located in the interchromatin compartment (IC)(blue), H3K27me3 marks are distributed within neighboring chromatin domain clusters; RNA Pol II is mainly enriched in chromatin lining the IC (purple), but also extends into the IC, whereas it is largely excluded from densely compacted chromatin regions (brown and yellow). **(G, H)** 3D image analyses of 3D SIM stacks (n: number of nuclei) shows the relative fraction of SC35 (red) and H3K27me3 (green) signals (G), and of active RNA Pol II (green) (H) in comparison to DAPI intensity classes 1-7 marked as black triangles. Complete image stacks from nuclei shown in A-F, marker distribution on DAPI intensity classes on individual nuclei and additional image stacks from two independent (replica) experimental series are provided in https://cloud.bio.lmu.de/index.php/s/rZxxkgYExonWLgy?path=%2FFig4

**Fig. 5:**
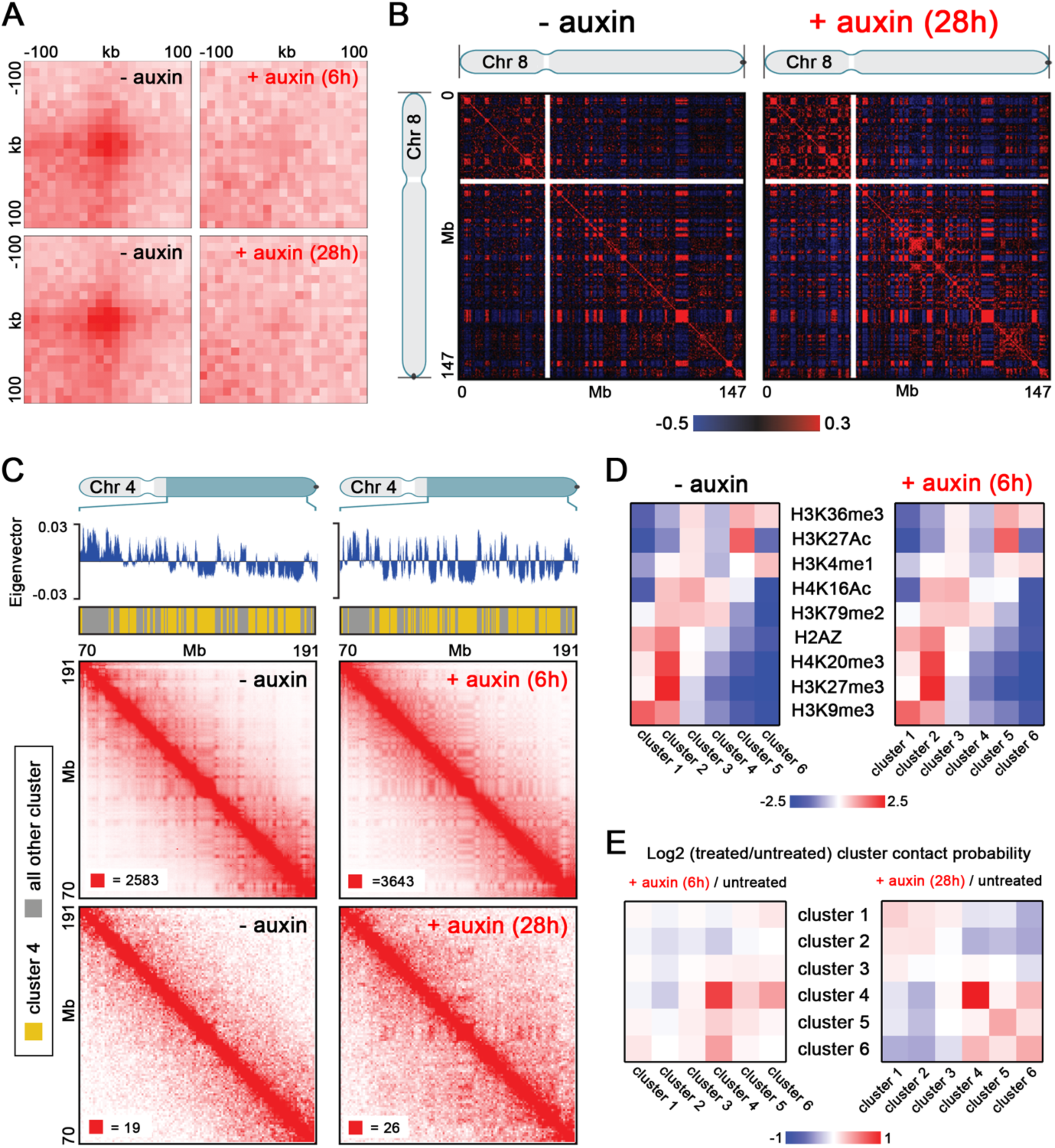
Hi-C data indicate elimination of chromatin loops, but maintenance of A and B compartments in cohesin depleted pre- and postmitotic cells. **(A)** Aggregate peak analysis (APA) plots using loops identified in HCT116-RAD21-mAC cells ^15^ before and after 6h of auxin treatment (top) or before and after 28h of auxin treatment (bottom). For each of the treated timepoints, the matched untreated control (harvested at the same time) is plotted next to it. The plot displays the total number of contacts that lie within the entire putative peak set at the center of the matrix. Loop strength is indicated by the extent of focal enrichment at the center of the plot. **(B)** Pearson’s correlation maps at 500 kb resolution for chromosome 8 before (left) and after (right) 28h of auxin treatment. The plaid pattern in the Pearson’s map, indicating compartmentalization, is preserved in cohesin depleted nuclei even after 28h of auxin treatment. **(C)** Contact matrices for chromosome 4 between 70 Mb and 191 Mb at 500 kb resolution before (left) and after (right) cohesin depletion. The 6h cohesin depletion time is shown on top, and 28h depletion time on the bottom. K-means clustering of histone modifications at 25 kb resolution into six clusters annotates loci corresponding to specific subcompartments. Interactions for loci in cluster 4 (arbitrary numbering, annotated in yellow on top tracks) are strengthened after both 6h and 28h of cohesin depletion. All loci belonging to clusters other than cluster 4 are annotated in gray in the top track. The max color threshold (red) of the heatmap is illustrated in the lower left corner of each heatmap, the minimum color threshold (white) is 0 reads. **(D)** Log-2 fold ratios of between-cluster Hi-C contact probabilities post- and pre-cohesin degradation are shown for the six clusters identified via K-means clustering of histone modifications. Cluster 4 shows strong contact probability enrichment after cohesin degradation at both the 6hr and 28hr timepoints. **(E)** For each of the 6 histone modification clusters, the average log2-fold enrichment for each histone modification over all loci in that cluster is shown both post- and pre-cohesin degradation. Patterns of histone modifications across the clusters as unchanged by cohesin degradation.

### Global features of higher order chromatin organization persist after cohesin depletion and are restored after mitosis in MLN despite the loss of loop domains

#### Maintenance and reconstitution of chromosome territories (CTs)

The capability of cohesin depleted cells to pass through an endomitosis prompted a careful comparison of the architecture of MLN compared with nuclei from control cultures and cohesin depleted cells on their way towards endomitosis (referred to as pre-mitotic cohesin depleted nuclei below). Maintenance of a territorial organization of interphase chromosomes in pre-mitotic, cohesin depleted cells and the reconstitution of CTs after endomitosis was demonstrated by chromosome painting of CTs 4, 12 and 19 (Fig. 2). In line with the near-diploid karyotype of HCT116 cells ^25^, two homologous territories of each painted chromosome were detected in interphase nuclei of both control (Fig. 2A) and pre-mitotic cohesin depleted cells fixed after 6h in auxin (Fig. 2B). Mitoses occurring in cohesin depleted cell cultures observed at this time revealed chromatid segregation, though frequently with misalignment (Fig. 2C). Most MLN fixed in cultures after 30h of auxin treatment revealed four painted territories for each delineated chromosome (Fig. 2D). Some MLN showed more than four painted regions with variable sizes, which were occasionally connected by thin chromatin bridges (Fig. 2D *right panel*, Supplementary Fig. 5). These observations may indicate that chromatids were torn apart by mechanic forces during lobe formation. Such disruptions might be enhanced, if we assume a higher level of relaxation and increased mechanical instability of chromosomes in cohesin depleted nuclei.

#### Evidence for co-aligned active and inactive nuclear compartments (ANC-INC) in cohesin depleted cell nuclei

Next, we tested the ability of cohesin depleted cells to preserve in addition to CTs other structural features of a compartmentalized nuclear architecture with active and inactive nuclear compartments described in the ANC-INC model ^19,20^. For this purpose, we compared DAPI-stained nuclei of cohesin depleted cells fixed after 6h in auxin, mostly comprising nuclei of the pre-mitotic interphase, and post-endomitotic MLN fixed after 30 h auxin treatment with control nuclei of cells cultured without auxin. Functionally relevant markers, delineated by immuno-detection, included SC35, an integral protein of splicing speckles involved in co-transcriptional splicing and transcriptional elongation ^26^, Ser5P-RNA Pol II, representing a transcription initiating form ^27^ (further referred to as RNA Pol II), and histone H3K27me3 conveying a repressed chromatin state ^28^. Two independent experiments (replicates 1 and 2) were performed with an interval of several months to test the long-term reproducibility of the results. 3D structured illumination microscopy (3D-SIM) was used to obtain stacks of nuclear serial sections from representative samples for further evaluation with our previously developed toolbox for 3D image analysis ^29^. This toolbox allowed highly resolved measurements of DNA intensity differences as proxies for chromatin compaction combined with the assignment of functional markers to regions of different compaction.

Figs. 3 and 4 present the combined results from replicates 1 and 2; for a separate presentation see Supplementary Fig. 6. Fig. 3A-C show typical mid-plane SIM sections of a control nucleus (A), a pre-mitotic cohesin depleted nucleus (B) and a post-endomitotic MLN (C). Color-coded voxels were attributed to seven intensity classes with equal intensity variance and represent the range of DAPI fluorescence intensities in 3D SIM nuclear serial sections. These color heat maps visualize local differences in DNA compaction ^29^. According to the ANC-INC model (see also Supplementary Table 1 for details of terminology), class 1 represents the interchromatin compartment (IC) with only sparse occurrence of DNA (blue). Chromatin domains (CDs) attributed to classes 2-7 form chromatin domain clusters (CDCs) with a nanoscale zonation of euchromatic and heterochromatic regions ^20,30^. Classes 2 and 3 (purple and red) comprise less compacted chromatin, including purple-coded chromatin directly bordering the IC, termed perichromatin region (PR). Classes 4-6 (orange, light brown, yellow) comprise facultative heterochromatin with higher compaction, class 7 (white) reflects the most densely compacted, constitutive heterochromatin. Enlargements of boxed areas in the three mid-plane nuclear sections of Fig. 3A-C exemplify chromatin domain clusters (CDCs) with a zonal organization of less compact chromatin domains at the periphery adjacent to the IC and higher compacted chromatin located in the CDC interior. Each CT is built from a number of CDCs, which in turn form higher order chromatin networks expanding throughout the nuclear space. 3D FISH with appropriate probes is required to identify individual CTs (Fig. 2) and CDCs (see Discussion).

**Fig. 6:**
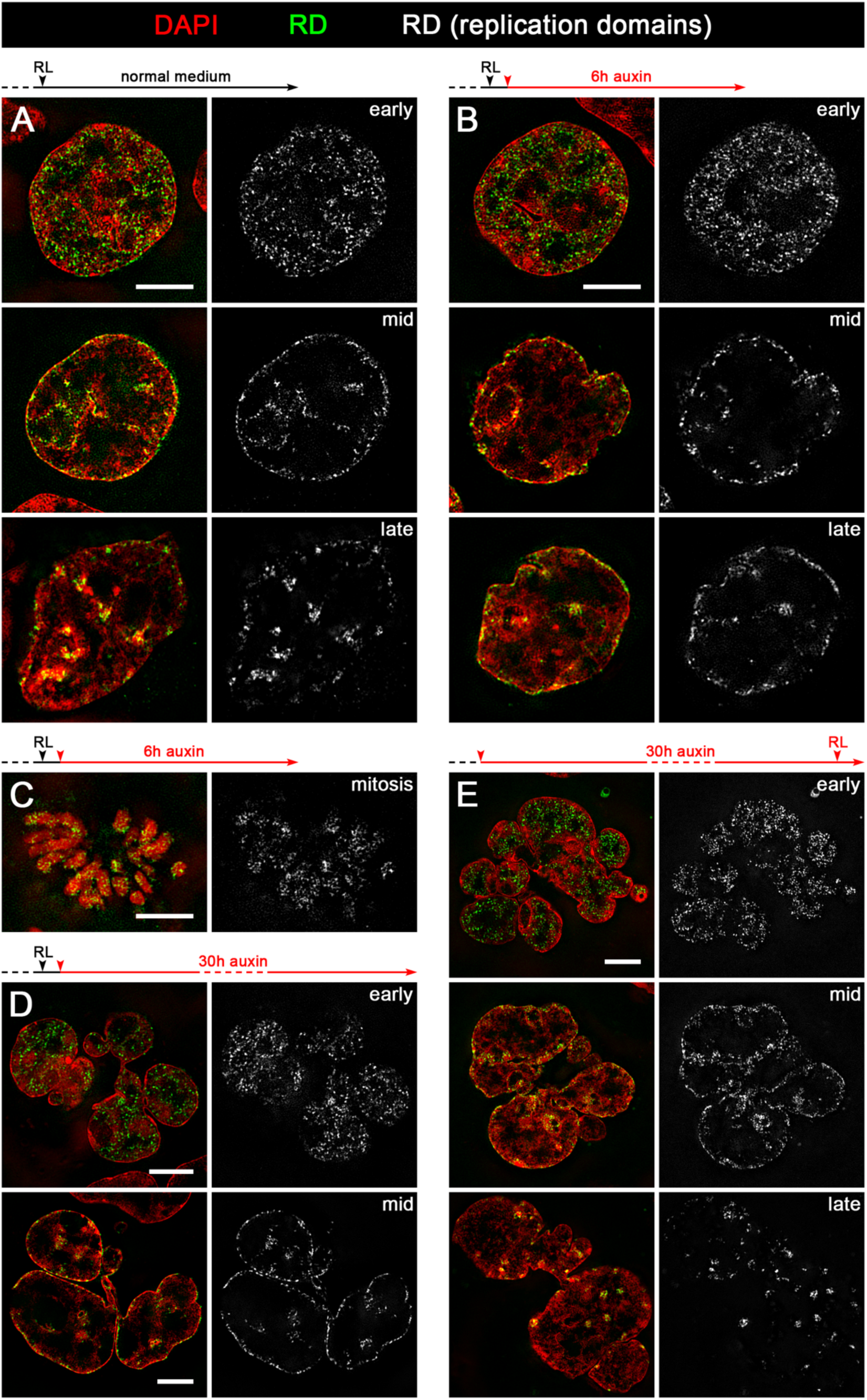
Maintenance, postmitotic rebuilding and de novo formation of typical replication patterns after cohesin depletion. **(A-E)** Overlay images (left) show representative SIM sections of DAPI stained nuclei (red) with replication domains (RDs)(green) identified by replication labeling (RL) in different stages of S-phase. RDs in the same nuclear sections are also displayed in gray (right). **(A)** Control nuclei fixed 6h after RL with typical patterns for early, mid and late replication, respectively. **(B)** Maintenance of the same typical replication patterns in nuclei of cohesin depleted, pre-mitotic cells fixed 6 h after RL.**(C)** Cohesin depleted mitotic cell with replication labeled chromatin domains obtained under conditions as described in (B). **(D)** RD patterns in individual lobuli demonstrate the ability of post-endomitotic MLN to restore RD patterns, generated by RL during the previous cell cycle. Cells were treated with auxin for 30h after RL. **(E)** RL carried out with MLN obtained after ∼30 h auxin treatment demonstrates *de novo* DNA synthesis with formation of new typical replication patterns. Scale bar: 5 µm.Raw data of complete image stacks from nuclei shown in A-E and additional image stacks from independent experimental series are provided in https://cloud.bio.lmu.de/index.php/s/rZxxkgYExonWLgy?path=%2FFig6

Relative fractions of voxels assigned to each of the seven DAPI intensity classes yielded similar patterns for control nuclei, pre-mitotic cohesin depleted nuclei and post-endomitotic MLN. (Fig. 3D). Fig. 3E presents estimates of nuclear volumes derived from 3D SIM serial sections. Whereas volumes of pre-mitotic cohesin depleted nuclei are similar to controls, the distinctly increased nuclear volume in MLN (30h auxin) corresponds with a further increase of a 2n DNA content immediately after endomitosis to a 4n DNA content (Supplementary Fig. 7) after passing through another round of DNA replication (see below). IC-channels expanding between lamina associated chromatin (Supplementary Fig. 8A-F) further illustrate the strikingly similar nuclear topography of higher order chromatin organization present in control nuclei, pre-mitotic cohesin depleted nuclei and post-endomitotic MLN. 3D image stacks reveal the integration of IC-channels and lacunas into an interconnected 3D network with direct connections to nuclear pores.^20,31^

**Fig. 7:**
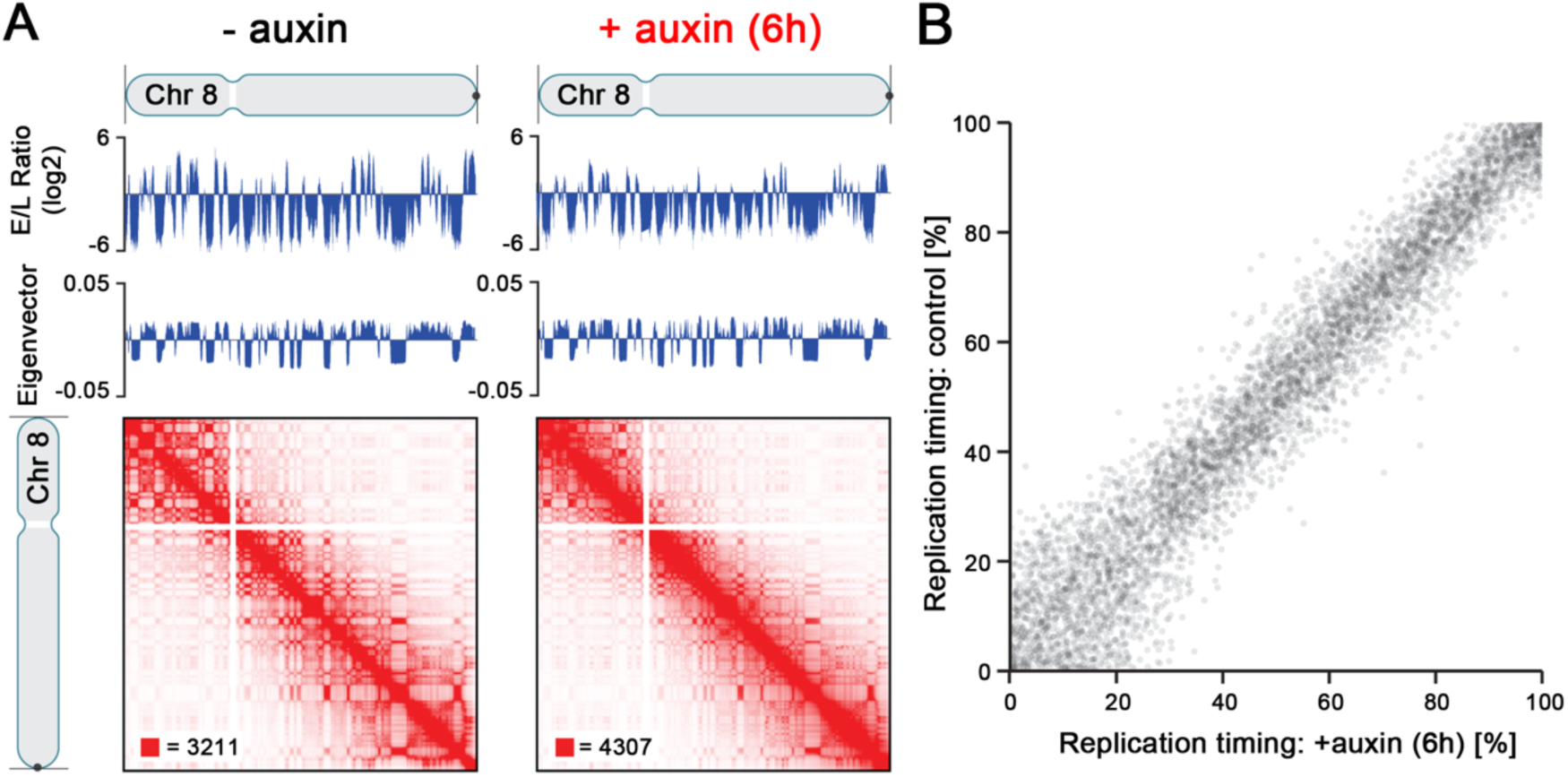
Hi-C and Repli-Seq data demonstrate the same replication timing for cohesin depleted and non-depleted control cells. **(A)** Contact matrices of chromosome 8 at 500 kb resolution along with the corresponding Repli-Seq early-to-late (E/L) ratio tracks at 50 kb resolution and the first eigenvectors of the Hi-C matrices corresponding to A/B compartmentalization. Replication timing along the genome is conserved, as shown by the correspondence of the untreated and auxin-treated Repli-Seq tracks. In addition, the correspondence between replication timing and genome compartmentalization (as indicated by the plaid pattern in the Hi-C map and the first eigenvector of the Hi-C matrices) is preserved after auxin treatment. **(B)** Scatter plot of replication timing (percentile of E/L ratio) in RAD21-mAC cells before (y-axis) and after (x-axis) auxin treatment.

**Fig. 8:**
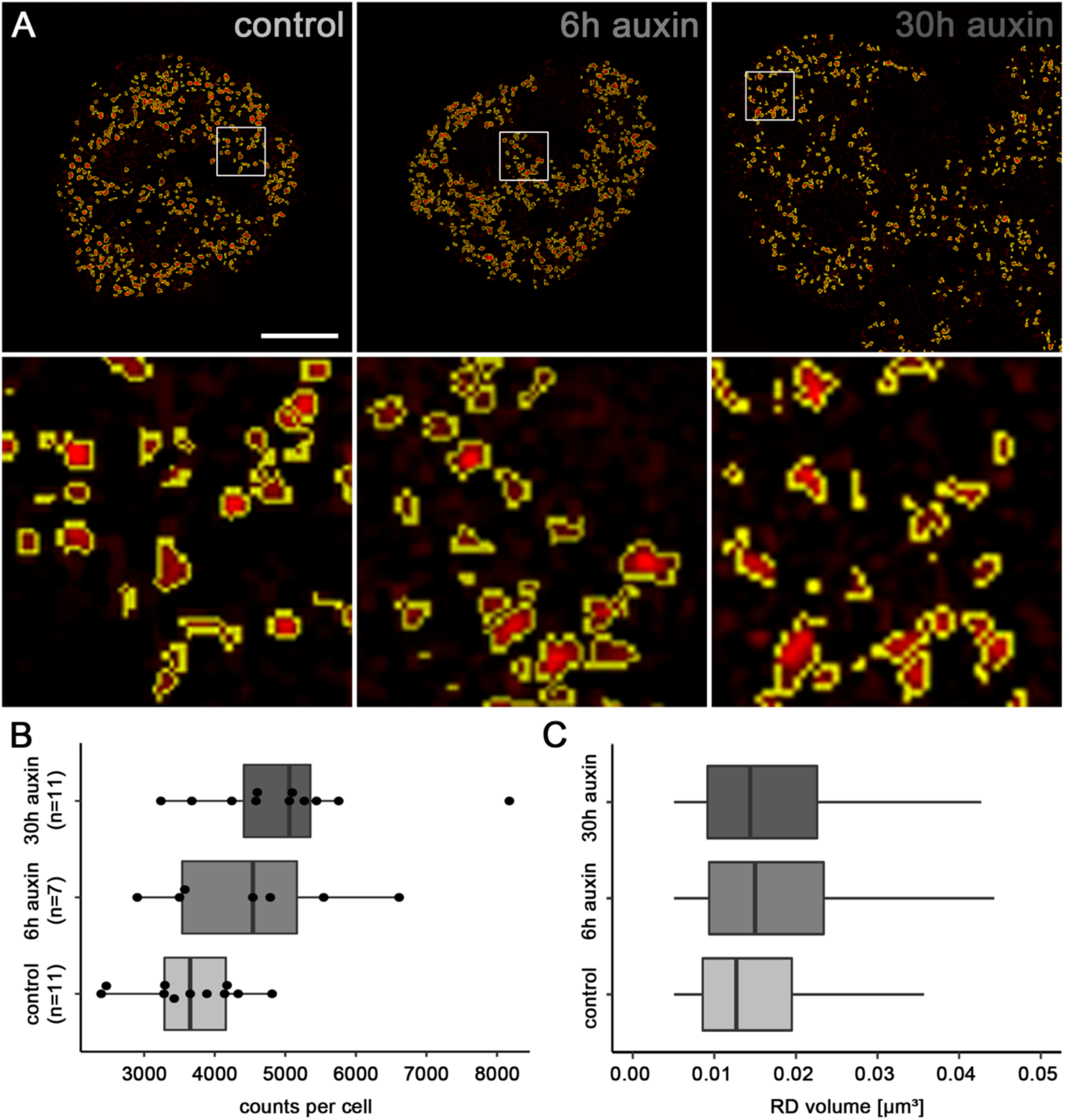
Segmentation of individual replication domains (RDs), pulse-labeled in early S-phase, indicate structural changes after cohesin depletion. **(A)** SIM nuclear mid-sections of nuclei with typical early S-phase patterns of RDs from a control culture (left) and an auxin treated culture (middle), both fixed 6 h after RL, and for 6h with auxin, and a multilobulated nucleus (right) *obtained* after 30h auxin treatment (compare Fig. 6 A-D). Enlargements of boxed areas show individual, segmented RDs displayed in red with segmented borders lined in yellow. Scale bar: 4 µm, 0.5 µm in inset magnifications. **(B)** Counts of segmented RDs plotted for 11 control nuclei, 7 cohesin depleted nuclei after 6h auxin, and 11 MLN after 30h auxin are presented as dots. Boxplots indicate the median with 25%-75% quartiles. **(C)** Boxplots with corresponding volume distributions of segmented, individual RDs (39.334 (control), 31.467 (6h auxin) und 55.153 (30h auxin). Lines demarcate minimum and maximum values. The non-parametric Mann-Whitney test revealed significant differences of RD counts between control nuclei and MLN (p = 0.012) and for RD volumes (p <0.0001 for control <-> 6h auxin, control <-> 30h auxin, and 6h auxin <-> 30h auxin) (Supplementary Table 3). Volumes of RDs with dimensions below the resolution limit of 3D-SIM (∼120 nm lateral / 300 nm axial) show the same size and were excluded from consideration. Accordingly, the lower limits of volumes between control nuclei and cohesin depleted nuclei are identical in contrast to the differences of the upper volume limits. Single values for individual nuclei are provided in https://cloud.bio.lmu.de/index.php/s/rZxxkgYExonWLgy?path=%2FFig3

Fig. 4A-F shows nuclear sections with DAPI stained DNA (gray) together with immunostained SC35 (red) and H3K27me3 (green) (A-C) or immunnostained RNA Pol II. Supplementary Fig. 6A-C demonstrates a range of compaction differences between SC35 marked speckles in both control and cohesin depleted nuclei. These examples illustrate the cell-to-cell variability of the nuclear landscape, which cannot be captured by a ‘typical’ one-for-all image. We did not further pursue the question, whether this structural variability reflects functional differences between individual cells in the non-synchronized cell populations studied here. In 3D SIM stacks of control and cohesin depleted nuclei we determined the relative fractions of voxels representing SC35, H3K27me3 and RNA Pol II, respectively, in relation to the seven DAPI intensity classes ^29^. By comparison of the relative fractions of marker voxels with DAPI related voxels, we tested for each class, whether a given marker showed a relative enrichment (over-representation) or relative depletion (under-representation) compared with the null-hypothesis of a random distribution (Fig. 4 G,H). Statistical tests are listed in Supplementary Table 3. Fig. 4G indicates a pronounced enrichment of SC35 in class 1 (IC), but a relative depletion in classes 2 and 3 (PR), and a virtual absence in higher classes. In contrast, H3K27me3, a marker of facultative heterochromatin, was under-represented in classes 1 and 2, but clearly enriched in classes 4 and 5. For RNA Pol II (Fig. 4H) we noted the most pronounced relative enrichment in class 2 and relative depletion in classes 4-7.

The separate presentation of both replicates (Supplementary Fig. 6D-E) consistently supports an enrichment of SC35 in class 1, and of H3K27me3 in class 4 and 5. The particular enrichment of H3K27me3 in classes 3 and 4 and depletion in class 7 is in line with its assignment as a marker for facultative heterochromatin ^32^. Enrichment-depletion patterns of RNA Pol II in the two replicates agree with respect to a general enrichment of RNA Pol II in the ANC (class 1-3), and a depletion within the INC, but differ markedly in quantitative details. Whereas replicate 1 shows a pronounced relative enrichment of this enzyme in class 1 and 2 in line with a relative depletion in classes 3 to 7, replicate 2 shows modest RNA Pol II enrichments in classes 2 and 3, together with relative depletions in classes 5-7, but unexpectedly also in class 1 (IC).

It is important to bear in mind that *relative* enrichments and depletions of epigenetic markers and functional proteins were defined in the 7 DAPI intensity classes. Differences between replicates 1 and 2 that represent snap-shots from the respective experiments may be attributed to unperceived differences of cell culture conditions. Notwithstanding these differences, both replicates support our major conclusion: Principal features of a compartmentalized organization with CTs and CDCs, pervaded by the IC in control nuclei were maintained in pre-mitotic, cohesin depleted nuclei and were rebuilt in post-endomitotic MLN, where individual macromolecules may penetrate into highly compacted CDs while macromolecular aggregates, such as a transcription machinery (RNA Pol II) or splicing machinery (SC35) may be excluded ^19,33^.

#### In situ Hi-C data indicate the maintenance/rebuilding of A and B compartments in cohesin depleted pre-mitotic nuclei and post-endomitotic MLN

In situ Hi-C of cell cultures, treated with auxin for 6 and 28h, respectively, prior to fixation, confirmed the disappearance of loop domains (Fig. 5A) in contrast to control cultures, whereas A and B compartments were maintained (Fig. 5B). Since most cells had passed an endomitosis with the formation of MLN after 28-30 h auxin treatment (Supplementary Fig. 4), we conclude that these findings are representative for both cohesin depleted pre-mitotic nuclei and post-endomitotic MLN. A heightened compartmentalization was noted in particular with regard to B-type chromatin, as previously described for pre-mitotic cohesin depleted cells ^15^. Strengthened interactions between this B-type compartment could be readily observed even in our low depth data from 28 h auxin treated cells (Fig. 5C, lower right panel, interactions between loci annotated in yellow). While the functional identity or significance of this particular B-type subcompartment remains unknown, we were able to identify by k-means clustering of histone modification data for HCT116-RAD21-mAC cells ^15^ a histone modification cluster (consisting of depletion of both activating marks like H3K36me3 and H3K27ac and repressive marks such as H3K27me3 and H3K9me3, but a mild enrichment of H3K79me2) that corresponded to the positions of this particular B-type subcompartment (Fig. 5D, E; cluster 4). Genome-wide analysis of the average Hi-C contact frequencies between the histone modification clusters demonstrated a strong enrichment for within-cluster contacts for this B-type subcompartment at both 6 h and 28 h after cohesin degradation, and additionally, at 28 h, mild cohesin-degradation induced enrichment of interactions between this B-type subcompartment and clusters enriched for repressive histone modifications as well as depletion of interactions with clusters enriched for activating histone modifications. The comparison of ensemble Hi-C data with microscopic data described above supports the argument that A/B compartments and ANC/INC compartments reflect the same structures (see Discussion).

#### Persistence of typical S-phase stage replication patterns after cohesin depletion

The next part of our study shows that the structural compartmentalization of pre-mitotic, cohesin depleted cells and post-endomitotic MLN corresponds with their functional capability to maintain replication domains and to proceed through S-phase. The temporal order of replication is highly coupled with genome architecture, resulting in typical patterns for early, mid and late replication timing ^34^. Replication domains (RDs) were chosen in our study as microscopically visible reference structures, which correspond to microscopically defined chromatin domains (CDs) and persist as stable chromatin entities throughout interphase and during subsequent cell cycles ^35-37^ (Supplementary Table 1). Replicating DNA was visualized by pulse replication labeling (RL) (see Methods). Control cultures were fixed 6h after RL (Fig. 6A), cultures prepared for cohesin depletion were further grown after RL for 1h under normal medium conditions and then exposed to auxin for 6h (Fig. 6 B,C) or 30h (Fig. 6D) before fixation. Both controls (A) and auxin-treated cells (B,D) revealed nuclei with typical RD patterns for different S-phase stages. This experiment demonstrates that different RD patterns persist during the subsequent pre-mitotic interphase of cohesin depleted cells (Fig. 6B) and can be fully reconstituted in post-endomitotic MLN (Fig. 6D). Notably, structural entities reflecting RDs pulse-labeled during S-phase can be identified along mitotic chromosomes (Fig. 6C). Fig. 6D demonstrates the ability of MLN to initiate a new S-phase with the formation of typical replication patterns.

#### Same replication timing for cohesin depleted and non-depleted control cells seen by Hi-C and Repli-Seq data

Using Repli-Seq and Hi-C analysis, replication timing was measured by the ratio of early to late replicating DNA and was found preserved upon cohesin depletion (Fig. 7A-B), consistent with a prior report ^38^. Additionally, the tight relationship between genome A/B compartmentalization and replication timing was similarly maintained in the absence of cohesin, exemplified for chr. 8 (Fig. 7A). Data were based on at least two replicates of each timepoint and confirmed reproducibility of results.

### Structural changes of replication domains in cohesin depleted nuclei

Finally, we tested whether cohesin depletion results in structural changes of individual RDs, detectable on the resolution level of 3D-SIM (Fig. 8). For this purpose, RD counts and RD volumes were evaluated in nuclei of three cultures: The ‘control’ culture was fixed 6h after RL together with the ‘6h auxin’ culture, which was incubated with auxin immediately after RL. The ‘30h auxin’ culture was fixed after 30h in auxin, when most cells had passed an endomitosis yielding a multilobulated cell nucleus. Nuclei with RD patterns typical for early S-phase at the time of pulse labeling were identified in the three fixed cultures and 3D serial image stacks of such nuclei were recorded with SIM and used for measurements in entire nuclei. It is important to note that an RD pattern generated by pulse labeling in a given nucleus is maintained after S-phase and after mitosis, independent of the time of fixation during the post-endomitotic interphase of MLN. Therefore, controls and auxin-depleted cells fixed 6 h after RL proceeded to G2, but still showed the early S-phase RD pattern. In the culture fixed 30 h in auxin, we identified MLN also showing early replication patterns. Fig. 8A shows examples of such nuclei from the control culture (left) and the cohesin depleted cultures fixed 6 h (middle) and from MLN cells fixed 30 h after RL (right). Fig. 8B presents average numbers of segmented RDs for individual nuclei. Fig.8C shows the results of volume estimates for individual RDs. Compared with controls, we noted an increase of both RD numbers and volumes together with an increase of heterogeneity (broader range of number and size distribution) in cohesin depleted pre-mitotic nuclei and post-endomitotic MLN. Based on the concordant increase of counts and volumes of segmented RDs in cohesin depleted nuclei in comparison with control nuclei, we tentatively conclude that cohesin is indispensable to prevent disintegration and decompaction of RDs (see Discussion).

### Effect of cohesin depletion on DNA halo induced chromatin loops

An effect of cohesin depletion on chromatin loop structure was supported by a DNA halo approach, a technique to investigate changes in chromatin organization at the level of DNA loops ^39^. Histone extraction in interphase nuclei by high-salt incubation triggers the extrusion of chromatin loops from a densely stained central chromatin core thus providing a measure of their size. DAPI stained nuclei of cohesin depleted cells (6h auxin treatment) exhibited halos that were significantly larger and more variable in shape in comparison to the defined and compacted halos of control cells (Supplementary Fig. 9) in line with the recently described observation that the cohesin-NIPBL complex compacts DNA by extruding DNA loops ^21^.

## Discussion

Our study demonstrates that multilobulated nuclei (MLN), that arise from cohesin depleted cells after passing through an endomitosis, retain the ability to rebuild a compartmentalized nuclear architecture. Whereas ensemble Hi-C confirmed the continued absence of chromatin loops and TADs in MLN as in pre-mitotic cohesin depleted nuclei, A and B compartments were fully restored in MLN in line with active and inactive nuclear compartments (ANC and INC, ^19,20^) revealed by 3D SIM. In light of the fundamental roles ascribed to cohesin, the capacity of MLN to initiate another round of DNA replication with stage-specific patterns of replication domains (RDs) was not expected.

Progression of cells into a disturbed and prolonged mitosis after cohesin depletion by Rad21 siRNA transfection was described in previous live cell studies covering ∼4h ^40^. By extending the live cell observation period up to 20h, we discovered a so far unreported endomitosis with chromatid segregation, but apparent failure to complete karyokinesis and cytokinesis. This failure may be attributed to the impact of cohesin for proper spindle pole formation and kinetochore-microtubule attachment (reviewed in ^7,8^). Notably, in vertebrates loading of cohesin onto DNA already occurs in telophase ^7^, which may be essential for subsequent cytokinesis and daughter cell formation. Factors promoting endomitosis and the formation of MLN are, however, complex and certainly diverse ^41^. Multipolar endomitosis with the formation of polyploid MLN occurs physiologically in megakaryocytes ^42^ and in (cohesin competent) tumor cell lines ^43^, in part entailing extensive chromosomal rearrangements ^44^. The observation of MLN as the mitotic outcome in ∼2% of HCT116-RAD21-mAC control cells exemplifies the spontaneous occurrence of MLN in a near-diploid tumor cell line.

Hi-C and related methods offer the great advantage of a genome wide approach to explore a nuclear compartmentalization at the DNA sequence level. This approach demonstrated a compartmentalized architecture of the landscape in cohesin depleted cell nuclei ^3,16^, but failed to detect the profound global morphological changes in post-endomitotic cohesin depleted MLN compared to cohesin depleted nuclei before passing through endomitosis. High-resolution microscopy is also the method of choice to examine the 3D structure of chromatin domain clusters (CDCs) with a zonal organization of repressed (condensed) and transcriptionally competent (decondensed) chromatin domains and the actual 3D configuration of the interchromatin compartment (IC) ^45^ with its supposed function as storage and transport system ^19^ that co-evolved with higher order chromatin organization ^46^. Our results exemplify the necessity to combine bottom-up with top-down approaches in ongoing 4D nucleome research, aimed at a comprehensive understanding of the structure-function relationships in complex biological systems.

We propose that microscopically defined ANC/INC compartments and A/B compartments, detected by ensemble Hi-C, represent the same functional compartments. Chromatin that contributes to the ANC and compartment A, respectively, is gene rich, transcriptionally active and typically located preferentially in the interior of mammalian cell nuclei, whereas both the INC and compartment B comprise gene poor, transcriptionally repressed chromatin of higher compaction, which is more prominent at the nuclear periphery (for review see ^20,47^). We further propose to equate microscopically defined chromatin domains (CDs) / RDs comprising several 100 kb (see below) that constitute functional building blocks of the ANC and INC with similarly sized compartment domains (see Supplementary Table 1) as functional building blocks of A and B compartments rather than with TADs ^48-50^. A correspondence of microscopically discernible RDs with TADs mapped by ensemble Hi-C has been favored in some studies ^49,50^

TADs represent genomic regions between several 100 kb up to >1 Mb in length, where DNA sequences physically interact with each other more frequently compared to sequences outside a given TAD ^47,51-53^. TADs, however, do not represent an individual chromatin structure, but a statistical feature of a cell population. Boundaries detected in Hi-C experiments are noted as transition points between TAD-triangles. They constrain, but do not restrict completely the operating range of regulatory sequences ^54^. Recently, super-resolution microscopy demonstrated the presence of TAD-like domains at the single-cell level ^55^. In cohesin depleted cells, a more stochastic placement of borders between TAD-like domains was detected ^56^. A role of IC-channels as additional structural boundaries between CDs and CDCs located on both sides, has been considered but not proven ^19^.

Early microscopic studies of the replicating genome during S-phase provided a first opportunity to explore its genome wide partitioning into discrete structural entities with a DNA content of ∼1 Mb, called replication domains (RD) or foci ^57,58^. We adopted the term ∼1 Mb chromatin domains in line with evidence that RDs persist as similarly sized stable chromatin units throughout interphase and during subsequent cell cycles ^35,36^. Later studies assigned an average DNA content of 400–800 kb to RDs/CDs ^18^, which can be optically resolved down to clusters of a few single replicons (150–200 kb) ^37,59^. Gene rich, early replicating domains form the A compartment, gene poor, later replicating domains the B compartment ^18^.

Our study confirms previous reports, which showed the maintenance of pulse-labeled RDs and the formation of S-phase specific replication patterns in cohesin depleted, pre-mitotic interphase cells ^30,38^. In addition, our study demonstrates the ability to re-constitute RDs in a typical pattern arrangement in post-endomitotic MLN. Moreover, MLN were able to initiate a new round of DNA replication with the formation of typical stage specific replication patterns under continued absence of cohesin.

These observations, however, do not imply that cohesin would be dispensable for RD structure. A comparison of numbers (counts) and volumes of individual RDs generated in early S-phase in nuclei of control cells and cells treated with auxin for 6 and 30 h respectively resulted in a significant increase both of RD numbers and RD volumes and also in a remarkably increased heterogeneity of these parameters after cohesin depletion. The near double amount of RD numbers in MLN (30h auxin) compared to controls was expected since MLN are generated as a result of an endomitosis with full separation of sister chromatids harboring RDs where labeled nucleotides were incorporated into both newly synthesized DNA strands in the previous cell cycle. In cohesin depleted cells treated with auxin after RL for 6h the increase of discernible RDs may result from an enhanced untethering of labeled sister chromatids compared to controls. At the time of fixation, both controls and cohesin depleted cells had likely reached the late S or G2-phase and labeled RDs had formed two separate sister chromatids within a given CT. Sister chromatids are kept together by cohesin at some sites, but are untethered at other sites and can dissociate from each up to few hundred nm^,61^. In cohesin depleted nuclei these untethered sites are likely increased. An increase of RD counts based on RD splitting should correspond with a decrease of RD volumes. Unexpectedly, we observed a remarkable volume increase in individual segmented RDs. This observation supports a role of cohesin in the compaction of chromatin structures exerted by chromatin loop extrusion 21 which could affect contact frequencies and thus explain at least in part the loss of TAD patterns in ensemble Hi-C experiments. Due to the resolution limit of 3D-SIM (∼120 nm lateral / 300 nm axial) these results must be viewed with caution: a fraction of RDs with sizes below this limit would show a putative size reflecting the diffraction limit, resulting in an overestimate of their volumes. To overcome these method-inherent limitations, imaging approaches with higher resolution, such as STORM/SMLM or STED are required to further clarify the influence of cohesin on RD structure ^60,61^. The increased heterogeneity of RDs volumes in cohesin depleted nuclei compared with controls, likely reflects the cell-to-cell shift of boundaries described for TAD-like domains in cohesin depleted cells ^56^. In summary, we tentatively conclude that cohesin plays an indispensable role for the structure of RDs/CDs but is dispensable for the formation of a compartmentalized nuclear organization. The current study may help to stimulate integrated research strategies with the goal to better understand the structure-function implications of the nuclear landscape.

New methods of super-resolved optical reconstruction of chromatin organization with oligopaints technology ^55^ or the combination of serial block-face scanning electron microscopy with in situ hybridization (3D-EMISH) ^62^ have opened up new ways to explore the geometrical variability of TAD-like structures in comparison with TADs identified by ensemble Hi-C and to close current gaps of knowledge on nuclear compartmentalization. Despite compelling evidence for chromatin loops, their actual 3D and 4D (space-time) organization is not known. Microscopic evidence for the formation of higher order chromatin arrangements based on nucleosome clutches or nanodomains ^30,55,56,63,64^ suggests that loops may be organized as much more compact structures with the potential implication that the diffusion of individual macromolecules into their interior may be constrained and the penetration of macromolecular aggregates is fully excluded ^33^. As a consequence, transcription and other nuclear functions may preferentially occur at the surface of chromatin clusters, dynamically remodeled to fulfill this condition. How dynamic changes of functionally defined higher order chromatin structures in space and time are related to changing functional requirements of cells at different levels of a hierarchical chromatin organization, defines major challenges for future studies. Such studies should also advance our still incomplete knowledge of cohesin functions.

### Materials & Methods

#### Cells and culture conditions

HCT116-RAD21-mAID-mClover cells (referred to as HCT116-RAD21-mAC cells in the manuscript) were generated and kindly provided by the Kanemaki lab (Mishima Shizuoka, Japan; ^14^). For a detailed description see Supplementary Fig. 1. Cells were cultured in McCoy’s 5A medium supplemented with 10% FBS, 2 mM L-glutamine, 100 U/ml penicillin, and 100 µg/ml streptomycin at 37°C in 5% CO_2_. For data shown in Supplementary Fig. 2B HCT116-RAD21-mAC cells and HCT-116 wild type cells were grown in in DMEM medium supplemented with 10% FCS, 2 mM L-glutamine, 50 μg/ml gentamicin. Cells were tested for mycoplasma contamination by confocal microscopy.

#### Auxin induced RAD21 proteolysis

Degradation of AID-tagged RAD21 was induced by addition of auxin (indole-3-acetic acid; IAA, Sigma Aldrich) to the medium at a final concentration of 500 µM (auxin stock solution 2 M in DMSO). In long term cultures fresh auxin-medium was added after ∼18h.

#### Immunodetection

Immunodetection of cohesin subunits RAD21, SMC1 and SMC3 was performed on cells grown to 80% confluency on high precision coverslips with respective antibodies all raised in rabbit (Abcam (RAD21), Bethyl laboratories (SMC1, SMC3)), detected with Cy3-conjugated goat anti rabbit antibodies (Dianova). Primary antibodies against SC35 (Sigma), RNA Pol II (Abcam) and H3K27me3 (Active Motif) were detected with either donkey anti-mouse Alexa 488 (Life technologies) or donkey-anti rabbit Alexa 594 (Life technologies). To meet the requirements for super-resolution microscopy with respect to an optimal signal-to-noise ratio and preservation of 3D chromatin structure, a protocol described in detail in ^65^ was followed. Cells were counterstained in 1 µg/ml DAPI and mounted in antifade mounting medium (Vectashield (Vector Laboratories)).

| **Antibodies** | **Source** | **Catalog number** |
| --- | --- | --- |
| RAD21 | Abcam | ab154769 |
| SMC1 | Bethyl laboratories | A300-055A |
| SMC3 | Bethyl laboratories | A300-060A |
| Goat anti rabbit Cy3 | Dianova | 111-165-045 |
| Mouse anti SC35 | Sigma | S4045 |
| Mouse anti RNA Pol II Ser5P | Abcam | ab5408 |
| Rabbit anti H3K27me3 | Active Motif | 39155 |
| Donkey anti mouse Alexa 488 | Life technologies | A21202 |
| Donkey anti rabbit Alexa 594 | Life technologies | A21207 |

#### Replication pulse labeling (RL)

##### 1. by replication scratch labeling

Cells cultivated on high precision coverslips (thickness 0.170 mm) grown to 50-80% confluency were transferred into a dry empty tissue dish after draining off excess medium. 30 µl of the prewarmed labeling solution containing 20 µM Cy3-dUTP (homemade) or Alexa 594-5-dUTP (Life technologies) was evenly distributed over the coverslip. With the tip of a hypodermic needle parallel scratches at distances of ∼100 µm were quickly applied to the cell layer. Cells were incubated for 1 min in the incubator, then a few ml of pre-warmed medium was added to the dish. After 30min medium was exchanged to remove non-incorporated nucleotides (for details see ^36^). This procedure preserves the RAD21-mClover fluorescence after labeling.

##### 2. by incorporation of 5-Ethynyl-dU (EdU) and detection via „click chemistry”

This approach was used for RL in MLN after 30h auxin treatment (compare Fig. 6E) since these cells are prone to detachment upon scratching. EdU was added at a final concentration of 10 µM to the medium for 15min. Incorporated EdU was detected according to manufactures instructions (baseclick) by a Cu(I) catalyzed cycloaddition reaction that covalently attaches a fluorescent dye containing a reactive azide group to the ethynyl-group of the nucleotide ^66^. For visualization of RDs, the dye 6-FAM-Azide (baseclick) at a final concentration of 20 µM was used.

After either labeling approach cells were washed in 1xPBS, fixed with 4% formaldehyde / PBS for 10 min and permeabilized with 0.5% Triton X-100/PBS/Tween 0.02% for 10 min. Cells were counterstained in 1 µg/ml DAPI and mounted in antifade mounting medium (Vectashield (Vector Laboratories); for details, see ^65^).

#### HI-C in situ analysis of untreated and auxin treated cells

HCT-116-RAD21-mAC cells were plated in 6-well plates with either complete media, or complete media with 500 µM auxin (IAA) for 6h (as in ^15^) or 28h (to enrich for post-mitotic cells with multilobulated nuclei). Cells were crosslinked with 1% formaldehyde directly on the plate for 10 minutes and then quenched with glycine. The crosslinked cells were then scraped off and *in situ* Hi-C was performed as in [14]. In brief, cells were permeabilized with nuclei intact, the DNA was digested overnight with MboI, the 5’-overhangs were filled in while incorporating bio-dUTP, and the resulting blunt end fragments were ligated together. Crosslinks were then reversed overnight, the DNA was sheared to 300-500 bp for Illumina sequencing, biotinylated ligation junctions were captured using streptavidin beads and then prepped for Illumina sequencing. We prepared 3 libraries (two biological replicates) each for each time point (untreated 6 hours, treated 6 hours, untreated 28 hours, treated 28 hours). All Hi-C data was processed using Juicer ^67,68^. The data was aligned against the hg19 reference genome. All contact matrices used for further analysis were KR-normalized with Juicer. Comparison of compartment strengthening to histone modification clusters was done as in ^15^. Histone modification data for 9 marks (H3K36me3, H3K27Ac, H3K4me1, H4K16Ac, H3K79me2, H2AZ, H4K20me3, H3K27me3, H3K9me3) generated from untreated and 6-hour treated cells in ^15^ was grouped into 6 clusters using k-means clustering. For the k-means clustering, the histone modification data was first converted into a z-score value for each mark in order to account for differences in the dynamic range between marks.

#### Repli-Seq of untreated or auxin-treated cells

HCT116-RAD21-mAC cells were synchronized in G1 with lovastatin as previously described ^69^. Briefly, cells were incubated with 20 µM Lovastatin (Mevinolin) (LKT Laboratories M1687) for 24 hours to synchronize in G1. 500 µM auxin or DMSO was added 6 hours before release from lovastatin block. To release from G1 block, lovastatin was washed away with 3 washes of PBS and warm media plus 2 mM Mevalonic acid (Sigma-Aldrich M4667) and 500 μM auxin or DMSO. Cells were released for 10, 14, 18, and 22 hours. 2 hours before the time point 100 μM BrdU was added to label nascent replication. After fixation, equal numbers of cells from each release time point were pooled together for early/late repli-seq processing ^17^. Repli-Seq data was processed as described in ^17^. In brief, data was aligned to the hg19 reference genome using bowtie2, deduplicated with samtools, and the log-2 ratio between early and late timepoints was calculated.

#### 3D DNA-FISH

Labeled chromosome painting probes delineating human chromosomes 4-(BIO), 12-(DIG) and 19-Cy3 were used. 30 ng of each labeled probe and a 20-fold excess of COT-1 DNA was dissolved per 1 µl hybridization mix (50% formamide/ 2xSSC/ 10% dextran sulfate).

Cells were fixed with 4% formaldehyde/PBS for 10 min. After a stepwise exchange with 0.5% Triton X-100/PBS, cells were permeabilized with 0.5% Triton X-100/PBS for 10 min. Further pretreatment steps included incubation in 20% glycerol (1h), several freezing/thawing steps in liquid N_2_, incubation in 0.1 N HCl (5 min) and subsequent storage in 50% formamide/2xSSC overnight. After simultaneous denaturation of probe and cells (2 min at 76°C), hybridization was performed at 37°C for 48h. After stringent washing in 0.1xSSC at 60°C, biotin was detected by streptavidin-Alexa 488 and DIG by a mouse-anti-DIG antibody conjugated to Cy5. Cells were counterstained in 1 µg/ml DAPI, and mounted in antifade mounting medium Vectashield (Vector Laboratories), (for a detailed protocol see ^22^).

#### DNA halo preparation

Cells were incubated for 6h in 500 µM auxin for cohesin depletion. DNA halo preparation was largely performed according to ^70^. After washing the cells in 1xPBS they were incubated for 10 min in a buffer at 4°C containing 10 mM Tris pH 8, 3 mM MgCl_2_, 0.1 M NaCl, 0.3 M sucrose, protease inhibitors (freshly added to the buffer prior to use) 1 μM pepstatin A, 10 μM E64, 1 mM AEBSF and 0.5% Nonidet P40. All the following procedures were performed at room temperature. Subsequently, DNA was stained for 4 min with 2 μg/ml DAPI. After 1 min in a second extraction buffer (25 mM Tris pH 8, 0.5 M NaCl, 0.2 mM MgCl_2_; protease inhibitors as in nuclei buffer and 1 mM PMSF were added fresh prior to use), cells were incubated 4 min in halo buffer (10 mM Tris pH 8, 2 M NaCl, 10 mM EDTA; protease inhibitors as in nuclei buffer and 1 mM DTT were added fresh prior to use). Finally, cells were washed 1 min each in two washing buffers (25 mM Tris pH 8, 0.2 mM MgCl_2_; the first buffer with and the second without 0.2 M NaCl). After 10 min fixation in 4% formaldehyde/PBS, cells were washed twice in 1xPBS and mounted on slides with Vectashield. Nuclear scaffolds and the faded DNA halos were imaged at a widefield microscope (Zeiss Axioplan 2, 100x/1.30 NA Plan-Neofluar Oil Ph3 objective; Axiovision software; AxioCam mRM camera). Both the total area (At) and the scaffold area (As) of each cell were manually segmented using the software Fiji and the DNA halo area (Ah) calculated as a subtraction of the two (Ah = At – As). The DNA halo radius was subsequently derived with the formula R = √(Ah/π). Four biological replicates were prepared and measured. For generation of plots and statistical analysis (Wilcoxon test) the software RStudio was used.

#### Confocal fluorescence microscopy

Confocal images were collected using a Leica SP8 confocal microscope equipped with a 405nm excitation laser and a white light laser in combination with an acousto-optical beam splitter (AOBS). The used confocal system has three different detectors, one photomultiplier tube (PMT) and two hybrid photodetectors (HyD). The microscope was controlled by software from Leica (Leica Application Suite X, ver. 3.5.2.18963). For excitation of DAPI, the 405 nm laser was used, for excitation of Alexa488, Cy3, STAR635P and Cy5, the white light laser was set to 499, 554, 633 and 649 nm, respectively. The emission signal of DAPI was collected by the PMT (412-512 nm), the emission signals of Alexa488 (506-558 nm), Cy3 (561-661 nm), STAR635P (640-750 nm) and Cy5 (656-780 nm) were collected by the two HyD detectors. Images were acquired with 42 nm pixel steps, 102 µs pixel dwell time and 2-fold line accumulation using a Leica HC PL APO 63x/1.30 NA Glycerol immersion objective. The frame size was 37 × 37 µm and the scan speed was 700 Hz. The size of the confocal pinhole was 1 A.U. Confocal image z-stacks were acquired with a step size of 330 nm.

#### Live cell microscopy for long term observations

For live cell imaging, cells were plated on poly-L-Lysine-coated glass bottom 2-well imaging slides (ibidi), allowing to image control and auxin-treated conditions in parallel. For DNA staining cells were incubated in media containing 500 nM SiR-DNA (Spirochrome) for 1h before imaging. Timelapse acquisitions were carried out on a Nikon TiE microscope equipped with a Yokogawa CSU-W1 spinning disk confocal unit (50 µm pinhole size), an Andor Borealis illumination unit, Andor ALC600 laser beam combiner (405 nm / 488 nm / 561 nm / 640 nm), and Andor IXON 888 Ultra EMCCD camera. The microscope was controlled by software from Nikon (NIS Elements, ver. 5.02.00). Cells were imaged in an environmental chamber maintained at 37°C with 5% CO2 (Oko Labs), using a Nikon PlanApo 60x/1.49 NA oil immersion objective and a Perfect Focus System (Nikon). Images were recorded every 15 min for 21h as z-stacks with two planes and a step size of 6 µm, unbinned and with a pixel size of 217 nm. For excitation of mClover and SiR-DNA, the 488 and 640 nm laser lines were used, respectively. Fiji software (ImageJ 1.51j) ^71^ was used to analyze images.

#### Quantitation of auxin induced RAD21-mAID-mClover degradation on single cells

1. *in fixed cells:* HCT-116-RAD21-mAC and HCT-116 wild type cells were treated with 500 μM auxin for 6h, fixed in 3.7% formaldehyde, permeabilized with 0.7% Triton X-100 for 15 min, counterstained with 1 μg/ml DAPI for 10 min and mounted in Vectashield mounting medium (Vector Laboratories). High-throughput imaging of single cells was performed at the wide-field microscope Operetta (40x/0.95 NA air objective; Harmony software; Jenoptik firecamj203 camera). The high-content images were analyzed on batch through a pipeline created with the Harmony software and nuclei identified based on DAPI signal. The nuclei found on the border of each field were removed and the remaining nuclei were selected based on morphology parameters, such as size and roundness. mClover intensities were then measured within the nuclear mask of the selected nuclei. The fluorescence intensities data were exported into tables and processed in RStudio to produce plots and statistical analysis. For each treatment, the measurements were combined from 3 biological experiments, each made of 2 technical replicates. mClover intensities measured from HCT-116 wild type cells were used as an estimate for the background level. A median of 10 A. U. (arbitrary units) was calculated for the nuclear mClover intensity in wild type cells (10.23 and 10.56 A. U. in the untreated and in the auxin treated wild type cells, respectively). This background value was subtracted from all values measured for the untreated and auxin-treated HCT-116-RAD21-mAC cells.
2. *in time lapse acquisitions*: Nikon spinning disk confocal live cell time lapses were acquired as described above. For the analysis the lower of the two planes showing interphase cells was used. The detailed description of segmentation and analysis scripts can be found as comments in the scripts which are deposited on GitHub (https://github.com/CALM-LMU/Cohesin_project.git). In brief, segmentation maps for nuclei in the SiR-DNA channel in confocal time lapses were obtained by a machine learning based pixel classification using Ilastik (standard settings). Segmentation maps were manually curated in order to analyze only individual nuclei. Nuclei were traced starting at time frame 1 until the cell entered mitosis and disappeared from the lower imaging plane. The generated segmentation maps were used to select single nuclei in the mClover channel. After background subtraction (modal gray value) the median intensity was measured for each labeled cell over time using Fiji. Only cells with a mClover intensity above 50 counts were included in the analysis. All data shown are normalized to their starting values. Cells surpassing a fluctuation above the 90 % quantile relative to their own rolling mean of 5 timepoints were filtered out. Plots were generated using Python.

#### DNA content assessment in individual nuclei by integrated DAPI intensity measurement

*DAPI stained nuclei* were acquired using the Nikon spinning disk system described above. Fixed samples of untreated control cells and cells treated with auxin for 30h were acquired as confocal image z-stacks in 35 planes with a step size of 300 nm using a Nikon PlanApo 100x/1.45 NA oil immersion objective. DAPI was excited with the 405 nm laser line. Segmentation and analysis scripts are described in detail in the scripts which are uploaded on GitHub (https://github.com/CALM-LMU/Cohesin_project.git). In brief, spinning disk confocal stacks of DAPI stained nuclei were used for a machine learning based pixel classification to obtain 3D segmentation maps of nuclei using Ilastik (standard settings). Segmentation maps were manually curated in order to analyze only individual non touching nuclei. After background subtraction (modal gray value) the integrated intensity was measured for each segmented DAPI stained nucleus by using Fiji. Plots were generated using R Studio.

#### Semi-automatic quantification of multilobulated nuclei (MLN) and mitoses

Image acquisitions were carried out on the Nikon spinning disk system described above. Using a Nikon PlanApo 100x/1.45 NA oil immersion objective tiled images (3×3 with 5% overlap and 131 nm pixel size) were acquired for each condition to increase the number of cells per field of view. Confocal image z-stacks were acquired in two planes with a step size of 6 µm in order to encompass cells, in particular mitotic cells, in different plane levels. DAPI and mClover were excited with 405 or 488 nm laser lines, respectively. All nuclei from each image (average 280 nuclei per image frame) were classified visually into morphologically normal nuclei, mitoses and multilobulated nuclei (MLN). In auxin treated cells nuclei with persistent RAD21-mClover fluorescence (∼2%) were excluded.

#### Structured illumination microscopy (SIM)

Super-resolution structured illumination imaging was performed on a DeltaVision OMX V3 system (Applied Precision Imaging/GE Healthcare) equipped with a 100x/1.4 NA UPlan S Apo oil immersion objective (Olympus), Cascade II:512 EMCCD cameras (Photometrics) and 405, 488 and 593 nm lasers (for detailed description, see ^72^). For sample acquisition oil with a refractive index of RI=1.512 was used. 3D image stacks were acquired with 15 raw images per plane (5 phases, 3 angles) and an axial distance of 125 nm and then computationally reconstructed (Wiener filter setting of 0.002, channel specific optical transfer functions (OTFs)) and color shift corrected using the SoftWoRx software (Applied Precision Imaging/GE Healthcare). After establishing 32-bit composite tiff stacks with a custom-made macro in Fiji/ImageJ2 (http://rsb.info.nih.gov/ij/), the data were subsequently aligned again to get a higher alignment precision. These images were then used for measurements in the Volocity software (Perkin Elmer).

#### Nuclear volume measurements

Volume measurements were done with the Volocity software (Version 6.1.2.). RGB image stacks were separated in their respective channels and then nuclei structures were obtained and segmented for volume measurements by using the following commands: 1. “Find Objects” (Threshold using: Automatic, Minimum object size: 200 µm^3^), 2. “Dilate” (number of iterations: 15), 3. “Fill Holes in Objects” and 4. “Erode” (number of iterations: 15). In ≈ 5% of cases we had to adjust these settings for the challenging task of nuclei segmentation. To confirm statistically significance of volume differences the Mann-Whitney test was applied.

#### Segmentation and quantification of replication domain (RD) signals

Aligned 3D SIM image stacks were used as RGB for object counting and volume measurements in the Volocity software. For each series between n=7 and n=11 nuclei were measured resulting in 31.000 – 55.000 single values for each series. Image stacks were separated into their respective channels. The segmentation of RD structures was performed with the following software commands: 1. “Find Objects” (Threshold using: Intensity, Lower: 32, Upper: 255), 2. “Separate Touching Objects” (Object size guide of 0,002 µm^3^) and 3. “Exclude Objects by Size”, excluding structures < 0,005 µm^3^. This cut-off level largely corresponds to the resolution limit of 3D-SIM (∼120 nm lateral / 300 nm axial). Exclusion of signals outside a selected nucleus was achieved by the commands “Intersect” and “Compartmentalize”. Segmentation of nuclei was realized by the following commands: 1. “Find Objects” (Threshold using: Intensity), 2. “Dilate”, 3. “Fill Holes in Objects” and 4. “Erode”. Measured values of individual object counts and segmented RD volumes were displayed as boxplots indicating the median with 25%-75% quartiles. To test for statistical significance a Mann-Whitney test was applied. R studio was used for generation of plots and statistical tests.

*3D assessment of DAPI intensity classes as proxy for chromatin compaction classification* Nuclei voxels were identified automatically from the DAPI channel intensities using Gaussian filtering and automatic threshold determination. For chromatin quantification a 3D mask was generated in ImageJ to define the nuclear space considered for the segmentation of DAPI signals into seven classes with equal intensity variance by a previously described in house algorithm ^29^, available on request. Briefly, a hidden Markov random field model classification was used, combining a finite Gaussian mixture model with a spatial model (Potts model), implemented in the statistics software R ^73,74^. This approach allows threshold-independent signal intensity classification at the voxel level, based on the intensity of an individual voxel. Color or gray value heat maps of the seven intensity classes in individual nuclei were performed in ImageJ.

#### Quantitative allocation of defined nuclear targets on 3D chromatin compaction classes

Individual voxels of fluorescent signals of the respective marker channels were segmented by a semi-automatic thresholding algorithm (accessible in VJ Schmid (2020). nucim: Nucleome Imaging Toolbox. R package version 1.0.9. https://bioimaginggroup.github.io/nucim/ XYZ-coordinates of segmented voxels were mapped to the seven DNA intensity classes. The relative frequency of intensity weighted signals mapped on each DAPI intensity class was used to calculate the relative distribution of signals over chromatin classes. For each studied nucleus the total number of voxels counted for each intensity class and the total number of voxels identified for the respective fluorescent signals for SC35, RNA Pol II, H3K27me3 was set to 1. As an estimate of over/under representations (relative depletion/enrichment) of marker signals in the respective intensity classes, we calculated the difference between the percentage points obtained for the fraction of voxels for a given DAPI intensity class and the corresponding fraction of voxels calculated for the respective signals. Calculations were performed on single cell level and average values over all nuclei used for evaluation and plotting. For a detailed description, see ^29^.

#### Statistics

Microscopic observations were verified from at least two independent series. For highly elaborate quantitative 3D analyses of super-resolved image stacks we selected between 4 and 39 nuclei for a given experiment with the only precondition of a high staining and structure-preserving quality. Data shown in Figs. 3, 4, 8, Supplementary Figs. 2 and 9 comprise merged data from different series, with links to data on individual experiments. Investigators were not blinded during the experiments and when assessing the outcome. Significance levels were tested by a non-parametric Wilcoxon test and a Bonferroni-Holm correction was used to avoid errors through multiple testing (see Supplementary_Table_3). The error bars represent the standard error of the mean. The variance was similar between the groups that were statistically compared.

## Data availability

Raw data used for Figs. 1-4, 6, 8, suppl. Figs. 2-7,9, additional „biological replicates” and complementary experiments can be accessed under https://cloud.bio.lmu.de/index.php/s/rZxxkgYExonWLgy.

Processed Hi-C and Repli-Seq data used in this study can be accessed under: https://www.dropbox.com/sh/y2w0xipwso9kgma/AAC2OihQJdIlrzqBBPX0zPcxa?dl=0 with GEO accession: GSE145099 and enter token cxmzqaeqzdefhsj into the box. Publicly available ChIP-Seq data used in this study are available at GEO accession: GSE104888.

## Acknowledgments

We thank Stefan Müller, University of Munich (LMU), for kindly providing labeled chromosome painting probes for 3D-FISH experiments and Irina Solovei, University of Munich (LMU), for generously providing antibodies, lab space and facilities to MC. We are most grateful to Toyoaki Natsume from the lab of Masato Kanemaki (Center of Frontier Research, National Institute of Genetics, Mishima, Shizuoka Japan) for providing HCT116-RAD21-mAC cells. KB was supported by the International Max Planck Research School for Molecular Life Sciences (IMPRS-LS). Microscopic images were acquired at microscopes of the Center for Advanced Light Microscopy (CALM) at the LMU Munich.

## Authors contributions

TC and ELA initiated the study; MC and TC conceived the microscopic experiments together with HH, KB and AM. KB, MC, AM and MGO performed experiments shown in Figs. 1-4,6,8 and Supplementary Figs. 2-7. AM and KB performed live cell and super-resolution/confocal microscopy; HH provided input on quantitative image analysis, including statistical analysis; AM performed segmentation analyses and VS 3D image analyses for chromatin density mapping data; MGO performed 3D rendering of nuclei. SM performed RAD21-mClover intensities by high-throughput imaging and DNA Halo experiments with support of MCC shown in Supplementary Figs. 2 and 9. Hi-C data were generated by SSPR and ELA with experimental support of NM (Fig. 5). Repli-Seq data (Fig. 7) were provided by DMG and KNK. HL provided input for the 3D imaging part and MCC for the replication part. MC and TC wrote the manuscript with support from all authors, in particular from ELA.

## Consent for publication

All authors read and approved the manuscript.

## Ethical approval and consent to participate

Not applicable

## Abbreviations

3D FISH: 3D fluorescence in situ hybridization
3D SIM: 3D structured illumination microscopy
AID: auxin inducible degron
ANC / INC: active / inactive nuclear compartment
CT: chromosome territory
CD(C): chromatin domain (cluster)
CTCF: CCCTC binding factor
DAPI: 4’,6-diamidino-2-phenylindole
EdU: 5-Ethynyl-2’-deoxyuridine
Hi-C: chromosome conformation capturing combined with deep sequencing
IC: interchromatin compartment
MLN: multilobulated nucleus
NC: nucleosome cluster
PBS: phosphate buffered saline
PBST: phosphate buffered saline with 0.02% Tween
PR: perichromatin region
RD: replication domain
RL: replication labeling
TAD: topologically associating domain

## References

1 Davidson, I. F. et al. DNA loop extrusion by human cohesin. Science 366, 1338–1345, doi: 10.1126/science.aaz3418 (2019).

2 Parelho, V. et al. Cohesins functionally associate with CTCF on mammalian chromosome arms. Cell 132, 422–433, doi: 10.1016/j.cell.2008.01.011 (2008).

3 Rao, S. S. et al. A 3D map of the human genome at kilobase resolution reveals principles of chromatin looping. Cell 159, 1665–1680, doi: 10.1016/j.cell.2014.11.021 (2014).

4 Zuin, J. et al. Cohesin and CTCF differentially affect chromatin architecture and gene expression in human cells. Proc Natl Acad Sci U S A 111, 996–1001, doi: 10.1073/pnas.1317788111 (2014).

5 Fudenberg, G. et al. Formation of Chromosomal Domains by Loop Extrusion. Cell Rep 15, 2038–2049, doi: 10.1016/j.celrep.2016.04.085 (2016).

6 Sanborn, A. L. et al. Chromatin extrusion explains key features of loop and domain formation in wild-type and engineered genomes. Proc Natl Acad Sci U S A 112, E6456–6465, doi: 10.1073/pnas.1518552112 (2015).

7 Jeppsson, K., Kanno, T., Shirahige, K. & Sjogren, C. The maintenance of chromosome structure: positioning and functioning of SMC complexes. Nat Rev Mol Cell Biol 15, 601–614, doi: 10.1038/nrm3857 (2014).

8 Mehta, G. D., Kumar, R., Srivastava, S. & Ghosh, S. K. Cohesin: functions beyond sister chromatid cohesion. FEBS Lett 587, 2299–2312, doi: 10.1016/j.febslet.2013.06.035 (2013).

9 Nishiyama, T. Cohesion and cohesin-dependent chromatin organization. Curr Opin Cell Biol 58, 8–14, doi: 10.1016/j.ceb.2018.11.006 (2019).

10 Peters, J. M., Tedeschi, A. & Schmitz, J. The cohesin complex and its roles in chromosome biology. Genes Dev 22, 3089–3114, doi: 10.1101/gad.1724308 (2008).

11 van Ruiten, M. S. & Rowland, B. D. SMC Complexes: Universal DNA Looping Machines with Distinct Regulators. Trends Genet 34, 477–487, doi: 10.1016/j.tig.2018.03.003 (2018).

12 Litwin, I., Pilarczyk, E. & Wysocki, R. The Emerging Role of Cohesin in the DNA Damage Response. Genes (Basel) 9, doi: 10.3390/genes9120581 (2018).

13 Merkenschlager, M. & Nora, E. P. CTCF and Cohesin in Genome Folding and Transcriptional Gene Regulation. Annu Rev Genomics Hum Genet 17, 17–43, doi: 10.1146/annurev-genom-083115-022339 (2016).

14 Natsume, T., Kiyomitsu, T., Saga, Y. & Kanemaki, M. T. Rapid Protein Depletion in Human Cells by Auxin-Inducible Degron Tagging with Short Homology Donors. Cell Rep 15, 210–218, doi: 10.1016/j.celrep.2016.03.001 (2016).

15 Rao, S. S. P. et al. Cohesin Loss Eliminates All Loop Domains. Cell 171, 305–320 e324, doi: 10.1016/j.cell.2017.09.026 (2017).

16 Haarhuis, J. H. & Rowland, B. D. Cohesin: building loops, but not compartments. EMBO J 36, 3549–3551, doi: 10.15252/embj.201798654 (2017).

17 Marchal, C. et al. Genome-wide analysis of replication timing by next-generation sequencing with E/L Repli-seq. Nat Protoc 13, 819–839, doi: 10.1038/nprot.2017.148 (2018).

18 Marchal, C., Sima, J. & Gilbert, D. M. Control of DNA replication timing in the 3D genome. Nat Rev Mol Cell Biol, doi: 10.1038/s41580-019-0162-y (2019).

19 Cremer, T. et al. The Interchromatin Compartment Participates in the Structural and Functional Organization of the Cell Nucleus. Bioessays 42, e1900132, doi: 10.1002/bies.201900132 (2020).

20 Cremer, T. et al. The 4D nucleome: Evidence for a dynamic nuclear landscape based on co-aligned active and inactive nuclear compartments. FEBS Lett 589, 2931–2943, doi: 10.1016/j.febslet.2015.05.037 (2015).

21 Kim, Y., Shi, Z., Zhang, H., Finkelstein, I. J. & Yu, H. Human cohesin compacts DNA by loop extrusion. Science 366, 1345–1349, doi: 10.1126/science.aaz4475 (2019).

22 Cremer, M. et al. Multicolor 3D fluorescence in situ hybridization for imaging interphase chromosomes. Methods Mol Biol 463, 205–239, doi: 10.1007/978-1-59745-406-3_15 (2008).

23 Jevtic, P., Edens, L. J., Vukovic, L. D. & Levy, D. L. Sizing and shaping the nucleus: mechanisms and significance. Curr Opin Cell Biol 28, 16–27, doi: 10.1016/j.ceb.2014.01.003 (2014).

24 Shu, Z., Row, S. & Deng, W. M. Endoreplication: The Good, the Bad, and the Ugly. Trends Cell Biol 28, 465–474, doi: 10.1016/j.tcb.2018.02.006 (2018).

25 Langer, S., Geigl, J. B., Ehnle, S., Gangnus, R. & Speicher, M. R. Live cell catapulting and recultivation does not change the karyotype of HCT116 tumor cells. Cancer Genet Cytogenet 161, 174–177, doi: 10.1016/j.cancergencyto.2005.01.013 (2005).

26 Lin, S., Coutinho-Mansfield, G., Wang, D., Pandit, S. & Fu, X. D. The splicing factor SC35 has an active role in transcriptional elongation. Nat Struct Mol Biol 15, 819–826, doi: nsmb.1461 [pii]10.1038/nsmb.1461 (2008).

27 Egloff, S. & Murphy, S. Cracking the RNA polymerase II CTD code. Trends Genet 24, 280–288, doi: S0168-9525(08)00128-5 [pii]10.1016/j.tig.2008.03.008 (2008).

28 Zhou, V. W., Goren, A. & Bernstein, B. E. Charting histone modifications and the functional organization of mammalian genomes. Nat Rev Genet 12, 7–18, doi: nrg2905 [pii]10.1038/nrg2905 (2011).

29 Schmid, V. J., Cremer, M. & Cremer, T. Quantitative analyses of the 3D nuclear landscape recorded with super-resolved fluorescence microscopy. Methods 123, 33–46, doi: 10.1016/j.ymeth.2017.03.013 (2017).

30 Miron, E. et al. Chromatin arranges in chains of mesoscale domains with nanoscale functional topography independent of cohesin. bioRxiv566638, doi:doi.org/10.1101/566638 (2020).

31 Schermelleh, L. et al. Subdiffraction multicolor imaging of the nuclear periphery with 3D structured illumination microscopy. Science 320, 1332–1336, doi: 10.1126/science.1156947 (2008).

32 Smeets, D. et al. Three-dimensional super-resolution microscopy of the inactive X chromosome territory reveals a collapse of its active nuclear compartment harboring distinct Xist RNA foci. Epigenetics Chromatin 7, 8, doi: 10.1186/1756-8935-7-8 (2014).

33 Maeshima, K. et al. The physical size of transcription factors is key to transcriptional regulation in chromatin domains. J Phys Condens Matter 27, 064116, doi: 10.1088/0953-8984/27/6/064116 (2015).

34 Dimitrova, D. S. & Berezney, R. The spatio-temporal organization of DNA replication sites is identical in primary, immortalized and transformed mammalian cells. J Cell Sci 115, 4037–4051 (2002).

35 Jackson, D. A. & Pombo, A. Replicon clusters are stable units of chromosome structure: evidence that nuclear organization contributes to the efficient activation and propagation of S phase in human cells. J Cell Biol 140, 1285–1295 (1998).

36 Schermelleh, L., Solovei, I., Zink, D. & Cremer, T. Two-color fluorescence labeling of early and mid-to-late replicating chromatin in living cells. Chromosome Res 9, 77–80 (2001).

37 Xiang, W. et al. Correlative live and super-resolution imaging reveals the dynamic structure of replication domains. J Cell Biol 217, 1973–1984, doi: 10.1083/jcb.201709074 (2018).

38 Oldach, P. & Nieduszynski, C. A. Cohesin-Mediated Genome Architecture Does Not Define DNA Replication Timing Domains. Genes (Basel) 10, doi: 10.3390/genes10030196 (2019).

39 Heng, H. H. et al. Chromatin loops are selectively anchored using scaffold/matrix-attachment regions. J Cell Sci 117, 999–1008, doi: 10.1242/jcs.00976 (2004).

40 Diaz-Martinez, L. A. et al. Cohesin is needed for bipolar mitosis in human cells. Cell Cycle 9, 1764–1773, doi: 10.4161/cc.9.9.11525 (2010).

41 Ovrebo, J. I. & Edgar, B. A. Polyploidy in tissue homeostasis and regeneration. Development 145, doi: 10.1242/dev.156034 (2018).

42 Mazzi, S., Lordier, L., Debili, N., Raslova, H. & Vainchenker, W. Megakaryocyte and polyploidization. Exp Hematol 57, 1–13, doi: 10.1016/j.exphem.2017.10.001 (2018).

43 Chen, J. et al. Polyploid Giant Cancer Cells (PGCCs): The Evil Roots of Cancer. Curr Cancer Drug Targets 19, 360–367, doi: 10.2174/1568009618666180703154233 (2019).

44 Joos, S. et al. Hodgkin’s lymphoma cell lines are characterized by frequent aberrations on chromosomes 2p and 9p including REL and JAK2. Int J Cancer 103, 489–495, doi: 10.1002/ijc.10845 (2003).

45 Rouquette, J., Cremer, C., Cremer, T. & Fakan, S. Functional nuclear architecture studied by microscopy: present and future. Int Rev Cell Mol Biol 282, 1–90, doi: 10.1016/S1937-6448(10)82001-5 (2010).

46 Cremer, T., Cremer, M. & Cremer, C. The 4D Nucleome: Genome Compartmentalization in an Evolutionary Context. Biochemistry (Mosc) 83, 313–325, doi: 10.1134/S000629791804003X (2018).

47 Rowley, M. J. & Corces, V. G. Organizational principles of 3D genome architecture. Nat Rev Genet 19, 789–800, doi: 10.1038/s41576-018-0060-8 (2018).

48 Zhao, P. A., Rivera-Mulia, J. C. & Gilbert, D. M. Replication Domains: Genome Compartmentalization into Functional Replication Units. Adv Exp Med Biol 1042, 229–257, doi: 10.1007/978-981-10-6955-0_11 (2017).

49 Pope, B. D. et al. Topologically associating domains are stable units of replication-timing regulation. Nature 515, 402–405, doi: 10.1038/nature13986 (2014).

50 Moindrot, B. et al. 3D chromatin conformation correlates with replication timing and is conserved in resting cells. Nucleic Acids Res 40, 9470–9481, doi: 10.1093/nar/gks736 (2012).

51 Lieberman-Aiden, E. et al. Comprehensive mapping of long-range interactions reveals folding principles of the human genome. Science 326, 289–293, doi: 10.1126/science.1181369 (2009).

52 Dixon, J. R. et al. Topological domains in mammalian genomes identified by analysis of chromatin interactions. Nature 485, 376–380, doi: 10.1038/nature11082 (2012).

53 Dixon, J. R., Gorkin, D. U. & Ren, B. Chromatin Domains: The Unit of Chromosome Organization. Mol Cell 62, 668–680, doi: 10.1016/j.molcel.2016.05.018 (2016).

54 Ibrahim, D. M. & Mundlos, S. The role of 3D chromatin domains in gene regulation: a multi-facetted view on genome organization. Curr Opin Genet Dev 61, 1–8, doi: 10.1016/j.gde.2020.02.015 (2020).

55 Boettiger, A. & Murphy, S. Advances in Chromatin Imaging at Kilobase-Scale Resolution. Trends Genet 36, 273–287, doi: 10.1016/j.tig.2019.12.010 (2020).

56 Bintu, B. et al. Super-resolution chromatin tracing reveals domains and cooperative interactions in single cells. Science 362, doi: 10.1126/science.aau1783 (2018).

57 Ma, H. et al. Spatial and temporal dynamics of DNA replication sites in mammalian cells. J Cell Biol 143, 1415–1425 (1998).

58 Nakamura, H., Morita, T. & Sato, C. Structural organizations of replicon domains during DNA synthetic phase in the mammalian nucleus. Exp Cell Res 165, 291–297, doi: 10.1016/0014-4827(86)90583-5 (1986).

59 Baddeley, D. et al. Measurement of replication structures at the nanometer scale using super-resolution light microscopy. Nucleic Acids Res 38, e8, doi: 10.1093/nar/gkp901 (2010).

60 Schermelleh, L., Heintzmann, R. & Leonhardt, H. A guide to super-resolution fluorescence microscopy. J Cell Biol 190, 165–175, doi:jcb.201002018 [pii]10.1083/jcb.201002018 (2010).

61 Cremer, C. & Masters, B. R. Resolution enhancement techniques in microscopy. Eur Phys J H 38, 281–344 (2013).

62 Trzaskoma, P. et al. Ultrastructural visualization of 3D chromatin folding using volume electron microscopy and DNA in situ hybridization. Nat Commun 11, 2120, doi: 10.1038/s41467-020-15987-2 (2020).

63 Otterstrom, J. et al. Super-resolution microscopy reveals how histone tail acetylation affects DNA compaction within nucleosomes in vivo. Nucleic Acids Res, doi: 10.1093/nar/gkz593 (2019).

64 Ricci, M. A., Manzo, C., Garcia-Parajo, M. F., Lakadamyali, M. & Cosma, M. P. Chromatin fibers are formed by heterogeneous groups of nucleosomes in vivo. Cell 160, 1145–1158, doi: 10.1016/j.cell.2015.01.054 (2015).

65 Markaki, Y., Smeets, D., Cremer, M. & Schermelleh, L. Fluorescence in situ hybridization applications for super-resolution 3D structured illumination microscopy. Methods Mol Biol 950, 43–64, doi: 10.1007/978-1-62703-137-0_4 (2013).

66 Salic, A. & Mitchison, T. J. A chemical method for fast and sensitive detection of DNA synthesis in vivo. Proc Natl Acad Sci U S A 105, 2415–2420, doi: 10.1073/pnas.0712168105 (2008).

67 Durand, N. C. et al. Juicebox Provides a Visualization System for Hi-C Contact Maps with Unlimited Zoom. Cell Syst 3, 99–101, doi: 10.1016/j.cels.2015.07.012 (2016).

68 Durand, N. C. et al. Juicer Provides a One-Click System for Analyzing Loop-Resolution Hi-C Experiments. Cell Syst 3, 95–98, doi: 10.1016/j.cels.2016.07.002 (2016).

69 Javanmoghadam-Kamrani, S. & Keyomarsi, K. Synchronization of the cell cycle using lovastatin. Cell Cycle 7, 2434–2440, doi: 10.4161/cc.6364 (2008).

70 Guillou, E. et al. Cohesin organizes chromatin loops at DNA replication factories. Genes Dev 24, 2812–2822, doi: 10.1101/gad.608210 (2010).

71 Schindelin, J. et al. Fiji: an open-source platform for biological-image analysis. Nat Methods 9, 676–682, doi: 10.1038/nmeth.2019 (2012).

72 Dobbie, I. M. et al. OMX: a new platform for multimodal, multichannel wide-field imaging. Cold Spring Harb Protoc 2011, 899–909, doi: 10.1101/pdb.top121 (2011).

73 Pau, G., Fuchs, F., Sklyar, O., Boutros, M. & Huber, W. EBImage--an R package for image processing with applications to cellular phenotypes. Bioinformatics 26, 979–981, doi: 10.1093/bioinformatics/btq046btq046 [pii] (2010).

74 R Core Team. (R Foundation for Statistical Computing, Vienna, Austria, 2013).

