## Supplementary Information for "Cohesin depleted cells rebuild functional nuclear compartments after endomitosis"

### Supplementary figures with legends

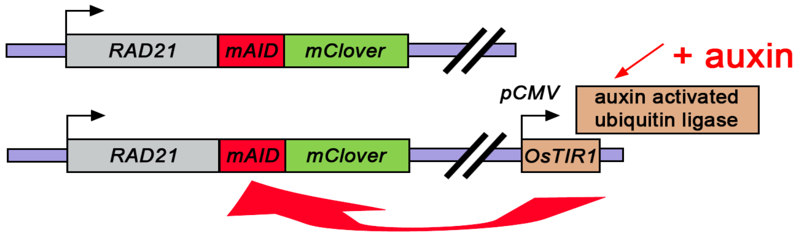


**Supplementary Fig. 1: Scheme of auxin induced RAD21 proteolysis**

Schematic draft for conditional proteolysis of RAD21 in HCT116-RAD21-mAID-mClover cells [12] further referred to in the manuscript as HCT116-RAD21-mAC. Both endogenous RAD21 alleles are fused with an auxin–inducible degron (AID) and fluorescent (mClover) reporter. In addition, the plant specific *OsTIR1* gene is integrated under the CMV promoter. For degradation, an ubiquitin ligase complex containing the plant specific OsTIR1 protein is activated via the association of auxin, a plant specific hormone involved in protein degradation pathways. Activation of TIR1 leads to ubiquitylation of the RAD21-AID-mClover fusion protein (for detailed information, see [1]).

1 Natsume, T., Kiyomitsu, T., Saga, Y. & Kanemaki, M. T. Rapid Protein Depletion in Human Cells by Auxin-Inducible Degron Tagging with Short Homology Donors. *Cell Rep* **15**, 210-218, doi:10.1016/j.celrep.2016.03.001 (2016).

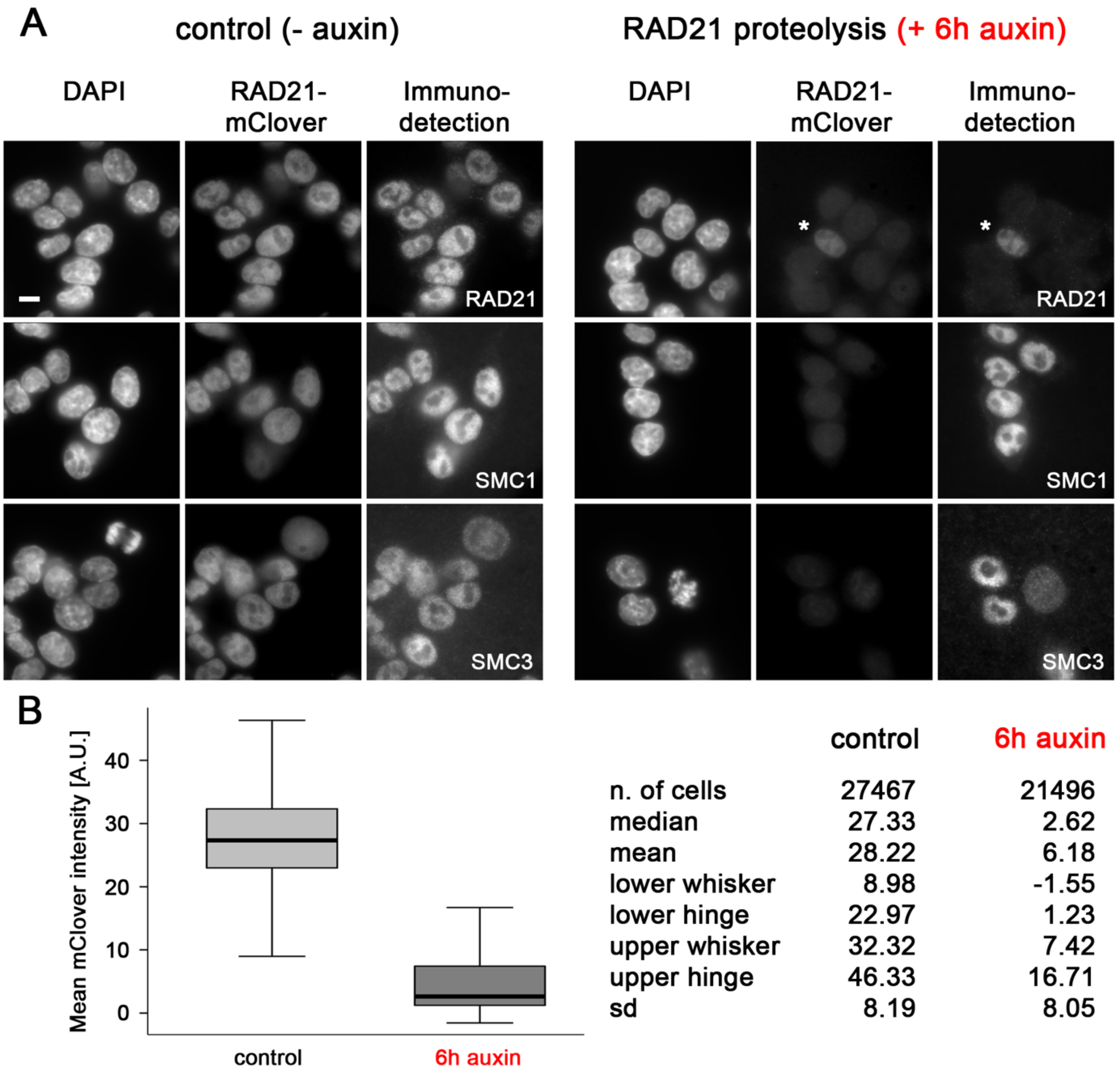


| | control | 6h auxin |
|---|---|---|
| n. of cells | 27467 | 21496 |
| median | 27.33 | 2.62 |
| mean | 28.22 | 6.18 |
| lower whisker | 8.98 | -1.55 |
| lower hinge | 22.97 | 1.23 |
| upper whisker | 32.32 | 7.42 |
| upper hinge | 46.33 | 16.71 |
| sd | 8.19 | 8.05 |

**Supplementary Fig. 2: RAD21-mClover proteolysis under auxin treatment**

**(A)** Immunodetection of the three major cohesin subunits RAD21, SMC1 and SMC3. Cells treated 6h with auxin (right panel) confirm the loss of RAD21 immunostaining in accordance with loss of RAD21-mClover fluorescence. Asterisk indicates a cell escaping RAD21 proteolysis. SMC1 and SMC3 immunostaining is maintained. Scale bar: 5 µm. **(B)** Averaged RAD21-mClover intensities recorded by high-throughput imaging from single cells of untreated controls (median = 27.33 A.U.) and auxin treated cells fixed after 6h in 500 µM auxin (median = 2.62 A.U.); p = < 0.001. For details, see Materials & Methods. The small overlap between cohorts is likely due to the small fraction of cells escaping AID induced RAD21 proteolysis in the auxin cohort and to cells that lack RAD21-mClover expression in control cells (compare Supplementary Fig. 3). Raw data and additional data are provided on https://cloud.bio.lmu.de/index.php/s/rZxxkgYExonWLgy?path=%2FSuppl_Fig2

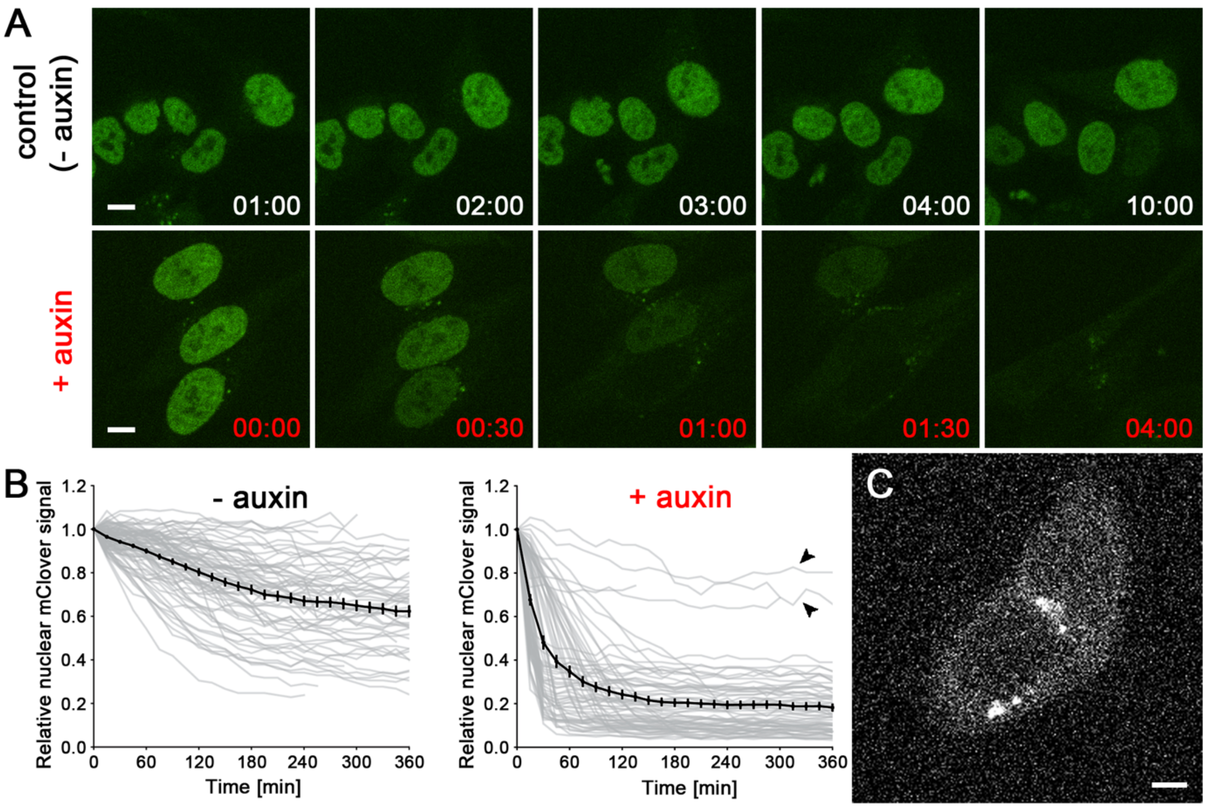


**Supplementary Fig. 3: Time course and quantitative measurement of auxin induced RAD21-mClover degradation based on single cell analyses from live cell observations**

**(A)** *Upper row:* selected time points of time lapse imaging (Δt=15min) in control cells show largely persistence of RAD21-mClover fluorescence over 10 h. *Lower row:* gradual decrease of RAD21-mClover fluorescence shown for selected time points in auxin treated cells recorded under same imaging conditions. RAD21-mClover fluorescence appears accomplished between time point 01:30-04:00 after addition of 500 μM auxin. Scale bar: 5 μm. Complete time lapse images (maximum projections) from data points shown in A and raw data of additional observations are provided in https://cloud.bio.lmu.de/index.php/s/rZxxkgYExonWLgy?path=%2FSuppl_Fig3. **(B)** Quantitative analysis of single cell nuclear RAD21-mClover fluorescence recorded over 6h from live cell experiment shown in (A). Time course of nuclear fluorescence was analyzed by use of automated image data analysis and segmentation tools (see materials and methods). In control cells nuclear fluorescence decreases to a mean value of ~60% of the starting value due to imaging related bleaching. Auxin treatment reduces RAD21-mClover fluorescence to an average of ~20% of the starting value, this value settles after ~3h. Arrows indicate cells that apparently escaped auxin degradation in this experiment. Control cells: n=82; auxin-treated cells: n=69. **(C)** Fluorescent degradation products of RAD21-mClover around the nucleus (cf. bright speckles) and in the cytoplasm, that can affect the results of the automated analysis.

**Note:** AID triggers proteasomal degradation of a tagged protein by ubiquitination, while its expression continues. It can be assumed that the remaining fluorescence in auxin treated cells mostly originates from RAD21-mClover in proteasomes associated with the cytoskeleton, centrosomes and the outer surface of the endoplasmic reticulum [2], while the fraction of RAD21 within an intact cohesin ring is neglectable. Note also the observation of a residual fluorescence of ~10% of mClover signals was described in the original publication by [1] after addition of 500 μM auxin.

1 Natsume, T., Kiyomitsu, T., Saga, Y. & Kanemaki, M. T. Rapid Protein Depletion in Human Cells by Auxin-Inducible Degron Tagging with Short Homology Donors. *Cell Rep* **15**, 210-218, doi:10.1016/j.celrep.2016.03.001 (2016).
2 Wojcik, C. & DeMartino, G. N. Intracellular localization of proteasomes. *Int J Biochem Cell Biol* **35**, 579-589, doi:10.1016/s1357-2725(02)00380-1 (2003).

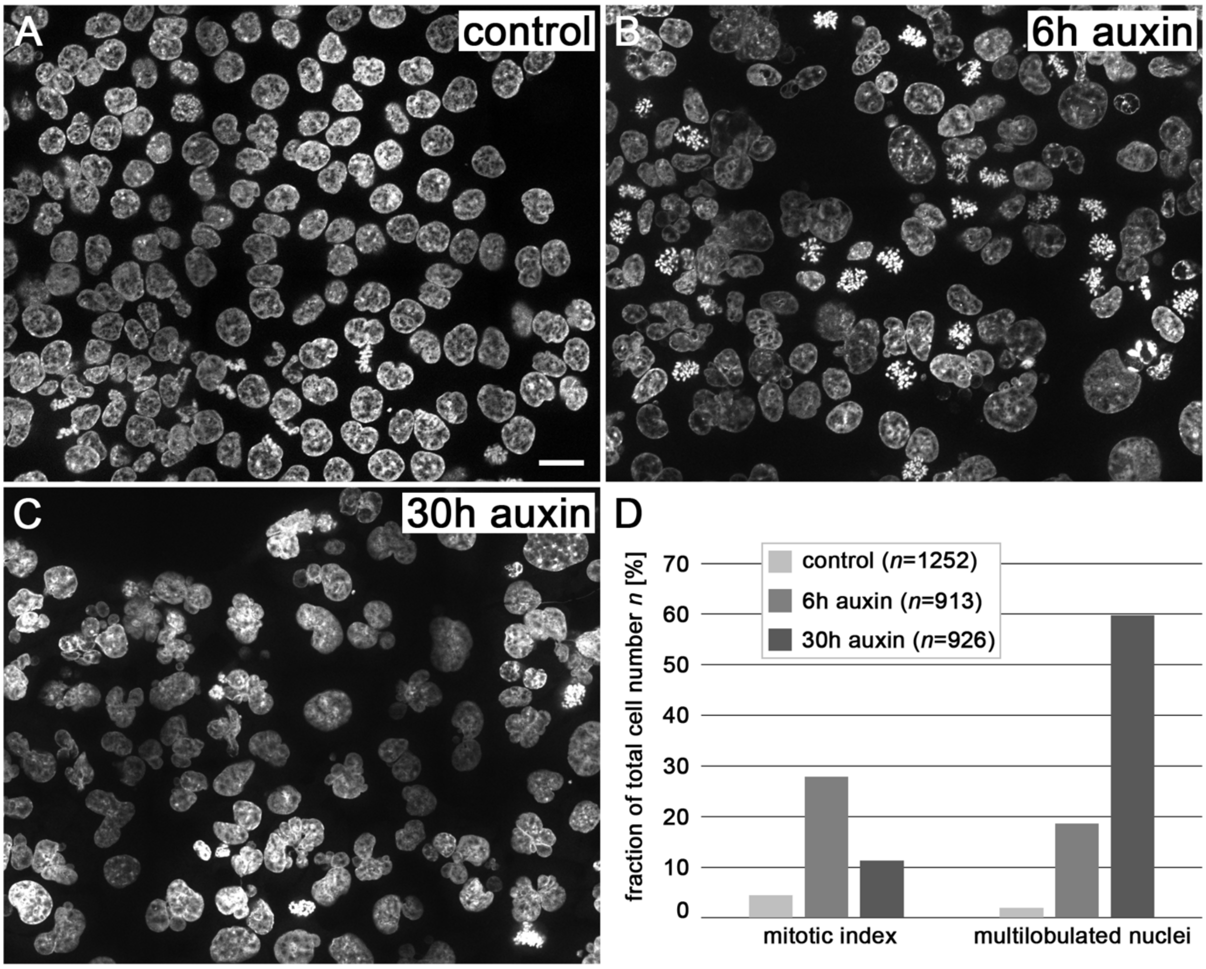


**Supplementary Fig. 4: Transient increase of mitotic index and fraction of MLN after cohesin depletion**

**(A-C)** Representative images (z-projections from two optical sections with $\Delta z$ = 6 µm) of **(A)** DAPI stained control nuclei; **(B)** nuclei fixed 6h after auxin treatment showing the accumulation of mitoses; **(C)** nuclei fixed 30h after auxin treatment with highly enriched MLN. Scale bar: 20 µm. **(D)** Quantification of mitotic index and fraction of MLN in control and cohesin depleted nuclei. Apoptotic and morphologically inconspicuous nuclei with distinct micronuclei were excluded from the MLN fraction. Additional observation fields are provided in

https://cloud.bio.lmu.de/index.php/s/rZxxkgYExonWLgy?path=%2FSuppl_Fig4

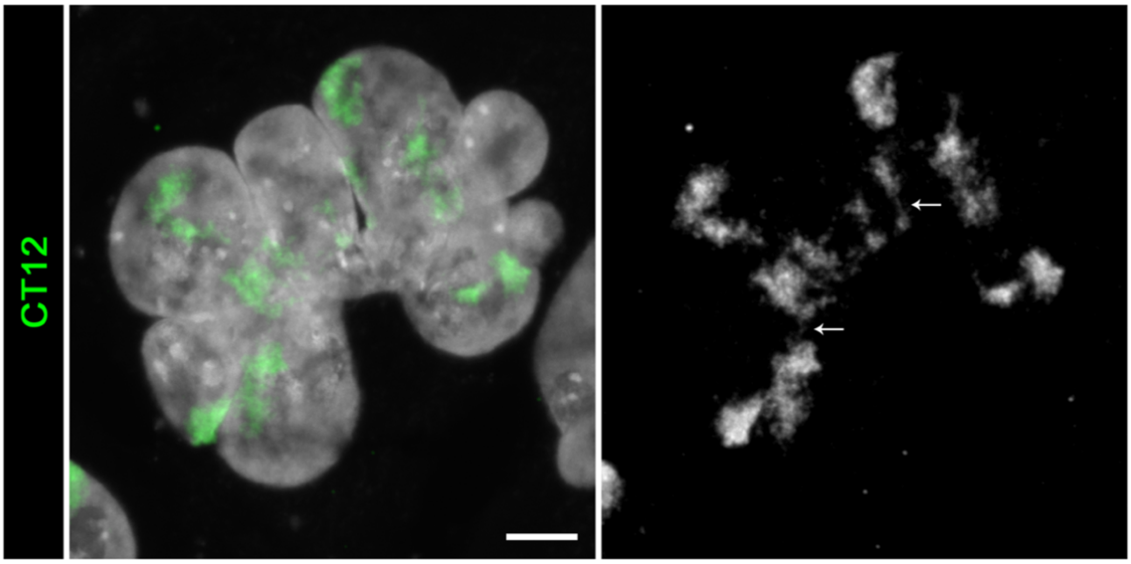


**Supplementary Fig. 5: Selective presentation of CT12 reveals a tearing apart of painted regions** *(Left)* Cohesin depleted postmitotic MLN (shown also in Fig. 2) with apparently >4 variably sized painted regions for CT12 in different lobuli (green). *(Right)* Selective presentation of painted regions (gray) reveal thin chromatin bridges between them (arrows). Scale bar: 5 µm

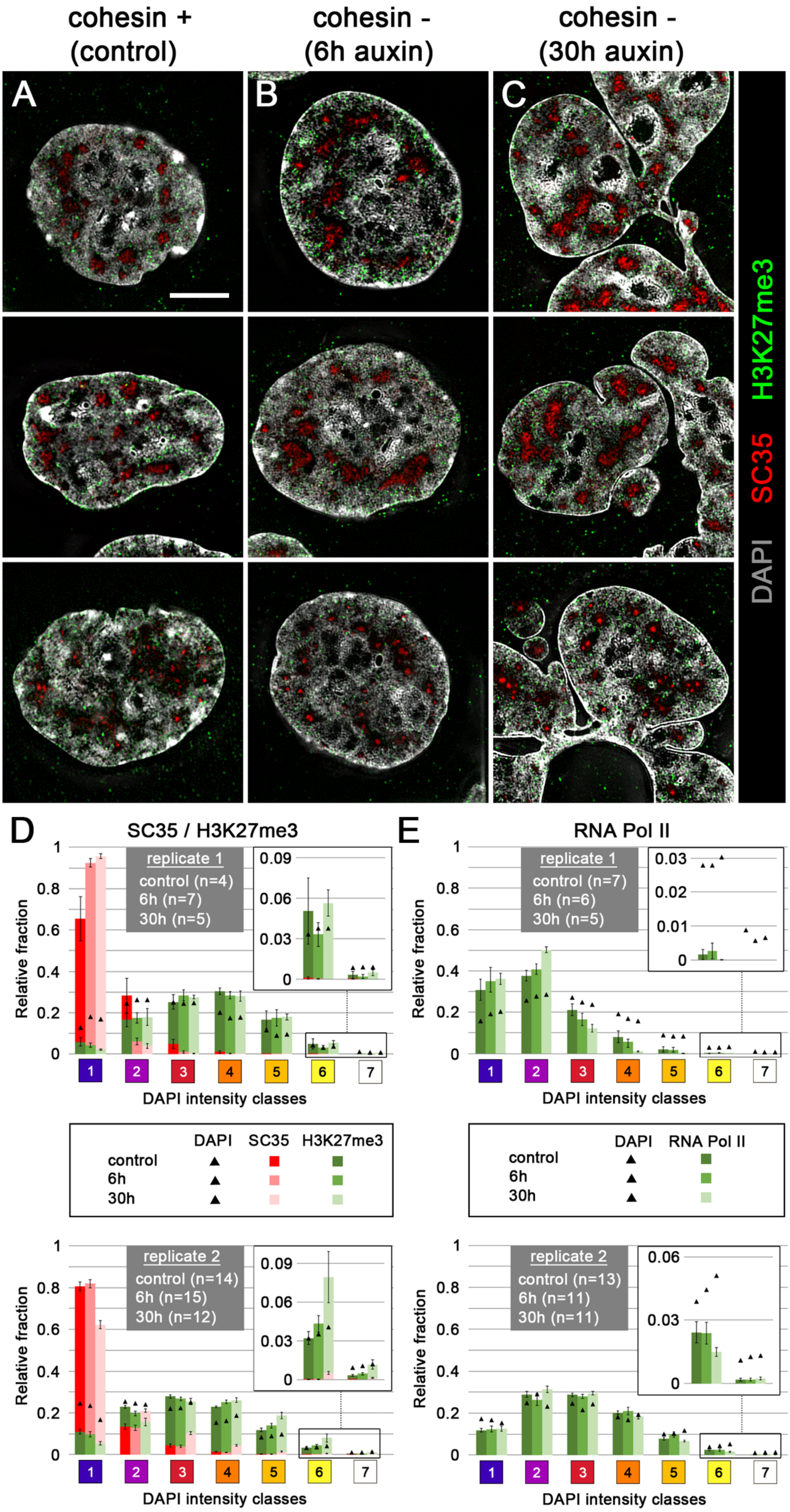

cohesin +
(control)
cohesin -
(6h auxin)
cohesin -
(30h auxin)
A
B
C
H3K27me3
SC35
DAPI
D
SC35 / H3K27me3
replicate 1
control (n=4)
6h (n=7)
30h (n=5)
Relative fraction
DAPI intensity classes
E
RNA Pol II
replicate 1
control (n=7)
6h (n=6)
30h (n=5)
DAPI SC35 H3K27me3
control
6h
30h
DAPI RNA Pol II
control
6h
30h
replicate 2
control (n=14)
6h (n=15)
30h (n=12)
replicate 2
control (n=13)
6h (n=11)
30h (n=11)

**Supplementary Fig. 6**: **Interexperimental variability of quantitative mapping of SC35, H3K27me3, RNA Pol II on chromatin compaction maps**

This Figure provides an extension of data shown in Fig. 4 to exemplify interexperimental variability. Replicates 1 and 2 were performed about one year apart from each other. (**A-C**) Mid-SIM sections from **(A)** control nuclei**, (B)** cohesin depleted nuclei fixed after 6 h auxin treatment**, (C)** partial sections of post-endomitotic MLN fixed 30 h after auxin treatment are shown with DAPI stained DNA (gray), immunostained SC35 (red) and H3K27me3 (green). SC35 marked speckles in IC lacunas demonstrate compaction differences between speckles in both control and cohesin depleted nuclei. (**D,E**) Separate presentation of replicates 1 and 2 with relative enrichments and depletions of SC35 and H3K4me3 (D) and RNA Pol II (E) in the 7 DAPI intensity classes (for explanation of details see the main text). Complete image stacks from nuclei shown in A-C and marker distribution on DAPI intensity classes in individual nuclei from the two different experiments are provided in

*https://cloud.bio.lmu.de/index.php/s/rZxxkgYExonWLgy?path=%2FSuppl_Fig6.*

**Note:**

Technical parameters for image recording and quantitative image analysis were kept constant in both experiments. Therefore, they are an unlikely source to explain interexperimental differences. Instead, unperceived biological differences between the cultures studied in the two experiments may explain why RNA Pol II was particularly enriched in classes 1 and 2 (IC and lining chromatin) in one experiment, and in classes 2 and 3 in another experiment. Although culture conditions appeared to be the same, parameters, which may affect the dynamics of higher order chromatin arrangements in cycling cells have remained elusive. The cultures were not synchronized and the two time points chosen for fixation of auxin treated cells (6h and 30h) have to be considered as snap-shots.

Different conformations of SC35 (shown in A-C) constituting a component in nuclear speckles composed of several protein complexes in a multilayered organization, was described [3], but underlying causes have remained elusive.

3 Fei, J. *et al.* Quantitative analysis of multilayer organization of proteins and RNA in nuclear speckles at super resolution. *J Cell Sci* **130**, 4180-4192, doi:10.1242/jcs.206854 (2017).

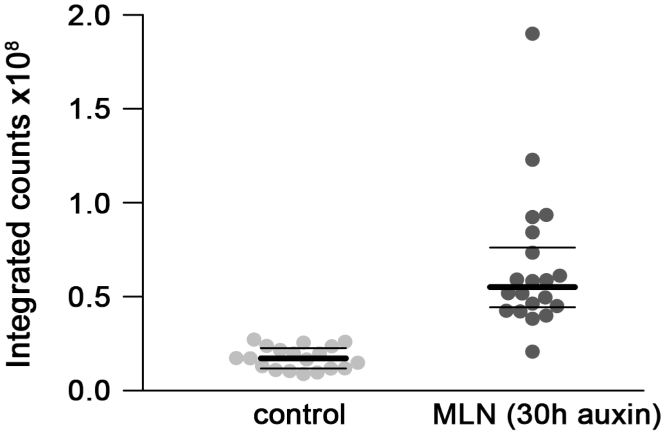


**Supplementary Fig. 7: DNA content measurements in controls and cohesin depleted postmitotic multilobulated nuclei (MLN)**

DNA content of single interphase nuclei based on integrated DAPI intensities of confocal sections in control cells (n=19) and postmitotic cohesin depleted MLN after 30h auxin treatment (n=20). Control cells show a narrow distribution reflecting the DNA content in G1, S, G2 phase (1n – 2n). MLN show an overall increased DNA content with a wide range. Note that these cells arise from an endomitosis with a 2n DNA content. MLN can pass through another full round of replication (compare Fig. 6E) increasing their DNA content up to 4n. Numerical data for each nucleus are provided in https://cloud.bio.lmu.de/index.php/s/rZxxkgYExonWLgy?path=%2FSuppl_Fig7

***Note:*** DAPI based single cell DNA content measurements may not reflect the absolute DNA content and a quantitative comparison of cells with highly different morphologies should be interpreted with caution [4]. However, the increased DNA content of cohesin depleted MLN is robust.


4 Roukos, V., Pegoraro, G., Voss, T. C. & Misteli, T. Cell cycle staging of individual cells by fluorescence microscopy. *Nat Protoc* **10**, 334-348, doi:10.1038/nprot.2015.016 (2015).

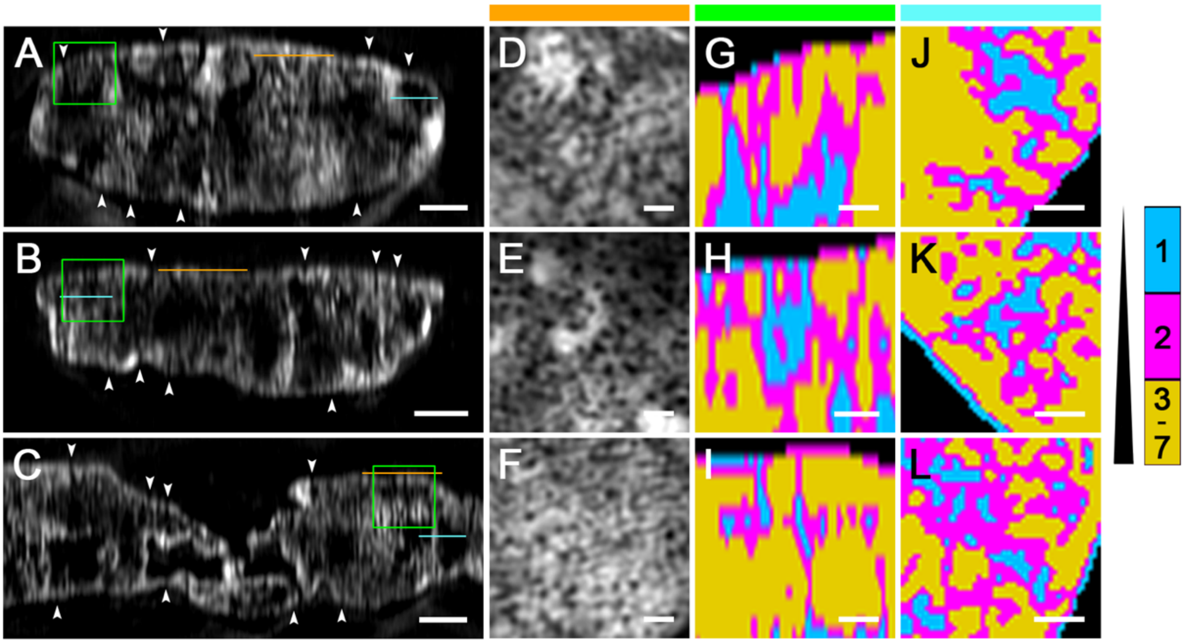


**Supplementary Fig. 8: Maintenance of a 3D network of the interchromatin compartment (IC) channel system after cohesin depletion and its reconstitution in MLN after mitosis**

**(A-C)** Z-sections from nuclei shown in Fig. 3A-C demonstrate IC-channels (arrowheads) extending between nuclear envelope associated chromatin domains into the nuclear interior, where they form wide IC-lacunas. Scale bars: 2 µm. **(D-F)** Apical XY-sections from respective nuclei (indicated as orange lines in A-C) delineating the passage of IC channels through peripheral heterochromatin at the nuclear lamina (not shown) where IC-channels are noted as black holes. Scale bars: 0.5 µm. **(G-I)** Inset magnifications from z-sections of nuclei shown in A-C (indicated as green frames) presented with color coded DAPI intensity classes 1 (cyan), 2 (magenta), 3-7 (merged in yellow; compare also Fig 3) further illustrate the extension of IC-channels from the nuclear periphery into the interior and the expansion of the IC into extended IC-lacunas (class 1) both within the control nucleus (G), and cohesin depleted pre- and postmitotic nuclei (H,I). The apparent predominance of a vertical channel alignment in z-sections is a consequence of the lower resolution in z (axial, ~250 nm) compared to xy (perpendicular to the optical axis, ~125 nm) [5]. **(I-L)** XY-sections from respective nuclei (indicated as cyan lines in A-C) demonstrate the occurrence of transversal channels from the lateral periphery towards the nuclear interior in line with a three-dimensional network. Scale bars: 0.5 µm. Complete image stacks are provided in https://cloud.bio.lmu.de/index.php/s/rZxxkgYExonWLgy?path=%2FSuppl_Fig8)

5 Schermelleh, L., Heintzmann, R. & Leonhardt, H. A guide to super-resolution fluorescence microscopy. *J Cell Biol* **190**, 165-175, doi:jcb.201002018 [pii]10.1083/jcb.201002018 (2010).

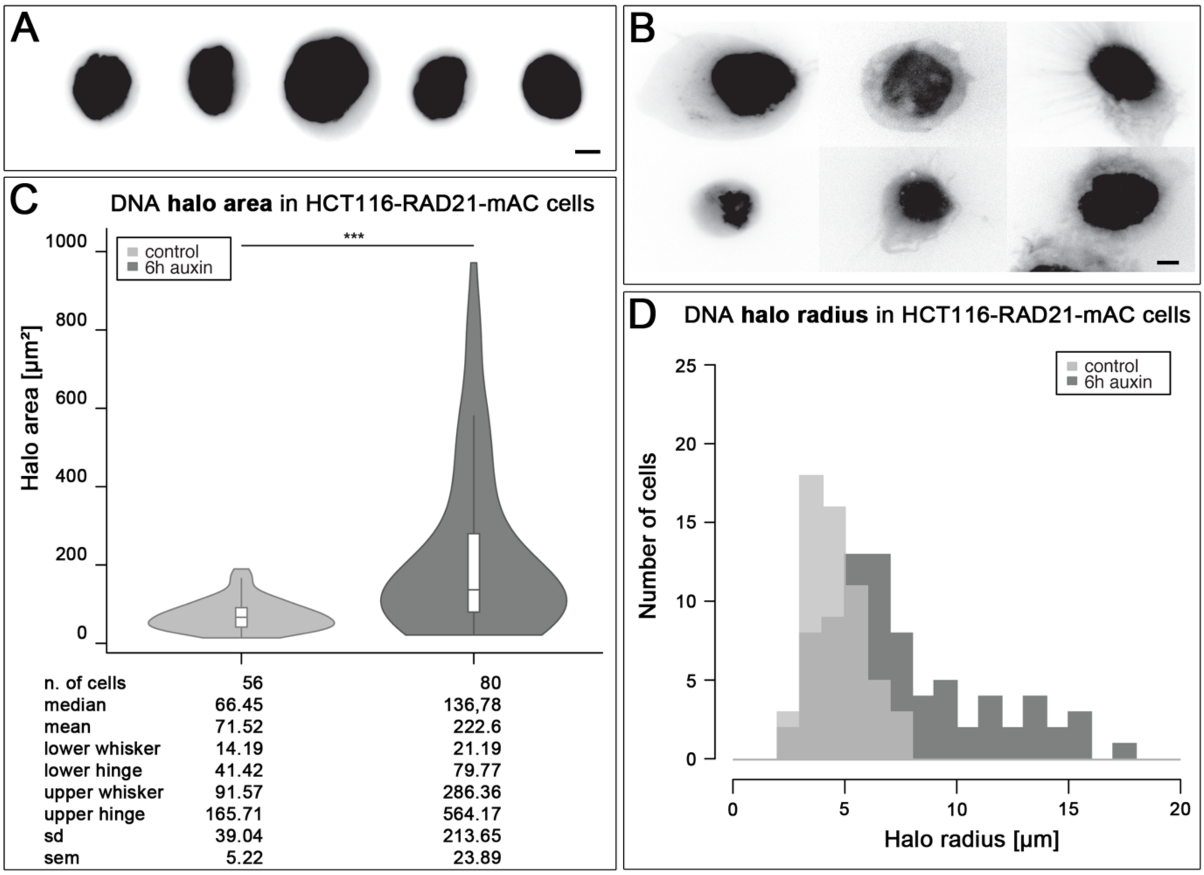


**Supplementary Fig. 9: Enlargement of DNA halos after cohesin depletion**

**(A-B)** Representative images of DNA halos stained with DAPI. The faded DNA halo surrounding a brighter insoluble nuclear scaffold corresponds to the DNA loops, whose extent reflects the degree of structural organization of chromatin. **(A)** Typical nuclei from control cells with small, rather uniform, and well delimited halos. **(B)** Halos of cohesin depleted cells show variable shapes and size, often ending up in extruded bundles of DNA fibers. Scale bar: 5 µm. **(C)** Violin plot showing the differences in area of the DNA halo between control and cohesin depleted cells (6h of auxin treatment), determined as described in Methods. Cohesin depleted cells show up to five times larger halos. p-value < 0.0001 (***) using Wilcoxon test. **(D)** Distribution of radial values of DNA halos for the two populations shown in (C). The radius was calculated as in the methods: $R = \sqrt{(Ah/\pi)}$. Data for replica are provided on https://cloud.bio.lmu.de/index.php/s/rZxxkgYExonWLgy?path=%2FSuppl_Fig9)

**Supplementary Table 1: Overview and explanatory notes on terminology for higher order chromatin structures.**

***Note:*** Microscopic and Hi-C studies have demonstrated different perspectives of the nuclear landscape in space and time. This table explains the author's use of the current terminology to describe the results of our present study together with a provisional attempt to point out relationships between specific terms used in microscopy and Hi-C, respectively. The current lack of a common terminology with which all investigators would agree, reflects unsolved gaps and inconsistencies in the present understanding of the functional hierarchy of chromatin and the functional nuclear organization as a whole (see question marks). Features detected by microscopy or Hi-C and related methods differ in important details but are believed to point to comparable structural and functional phenomena of the nuclear landscape, whose integration into a common model is an important future task.

| **microscopic methods** | **Hi-C and related methods** | **explanatory notes** |
|---|---|---|
| nucleosome clutch (NC) nanodomain (ND) [6-8] | ? | NCs/NDs with sizes of a few kb may represent the smallest structural entities in the hierarchy of chromatin organization above the level of individual nucleosomes. Nucleosomes in NCs/NDs may be so densely packed that macromolecular aggregates can exert their functions only at the surface of NCs/NDs. 4D organization of NCs/NDs has remained elusive. |
| chromatin loop[9-13], [14-16] | | Chromatin loops were described in microscopic and Hi-C studies. They are frequently anchored by a cohesin ring at a pair of convergent CTCF binding sites, but their 4D organization is still not well understood. The 4D organization of such loops in living cells has remained elusive. Current perspectives range from an open architecture, where DNA targets in the loop interior are easily accessible for macromolecular complexes to highly compact structures, which constrains the accessibility of individual macromolecules and excludes larger macromolecular complexes. |
| ? | compartmental domain [17] | Compartmental domains with a DNA content of ~10-15 kb were detected in *Drosophila* with high-resolution Hi-C. Here, a cohesin ring may tether two neighboring loops encompassing an active and a repressed compartmental domain, or a single loop spanning both an active and inactive compartmental domain. |
| TAD-like domain [18] | contact domain [15]<br><br>topologically associating domain (TAD) [16,19,20] | Contact domains with a median DNA content of 185 kb (range 40 kb to 3 Mb) were first described in human cell types. They reflect enhanced contact frequency among all loci in a contiguous interval and can form as a result of loop extrusion [21,22] or compartmentalization [20]. The term topologically associating domain (TADs) was introduced to define structural entities in the order of a few hundred kb to 1 Mb, but depending on the author, TAD is often used to refer to a wider size range [19,23]. Contact domains/TADs detected in population Hi-C studies do not represent an individual chromatin structure, but rather a statistical feature of a cell population. Super-resolution microscopy demonstrated the presence of TAD-like domains at the single-cell level. Though sharp domain boundaries were detected between neighboring TAD-like domains in single cells, cell-to-cell shifts of boundaries can abolish the detection of TADs in population Hi-C studies. |

| | | |
|---|---|---|
| replication domain (RD) [24-33]<br><br>~1 Mb chromatin domain (CD) [10,25,34,35] | A/B compartment domains [9,19,20,36] | RDs with an estimated DNA content of ~1Mb were first detected in mammalian cells by pulse-labelling of replicating DNA during S-phase with halogenated nucleotides. At high resolution, RDs represent assemblies of several replicons with ~100–200 kb. RDs are stably maintained over subsequent cell cycles independent of their association with the transcription machinery. Chromatin assembled in early replicating RDs is enriched with active genes, whereas mid- and late-replicating RDs comprise mostly repressed chromatin. The definition of ~1Mb chromatin domains (CDs) was based on replication domains (RDs) in early microscopic studies. CDs with less compact chromatin are enriched with epigenetic marks of transcriptionally competent chromatin, whereas CDs with chromatin of higher compaction are enriched with marks for repressed chromatin. We suggest to equate active and repressed CDs/RDs with active (A) and repressed (B) compartment domains, respectively, which altogether represent the global nuclear compartmentalization in A and B compartments (see below), insofar as they correlate with compartment boundaries rather than extrusion-mediated boundaries. Despite the evidence for active and repressed CDs, the observation of compartmental domains, where both active and inactive segments are present within a given contact domain (see above) argues that CDs may not be structurally and functionally uniform entities. An active gene, for example, located within a repressed CD provides an anomaly like a corn of pepper in a sugar box. More detailed comparisons between the nuclear landscapes present in different cell types and species are required to close this important gap of knowledge. |
| chromatin domain cluster (CDC) [34]<br>CD chains [6] | ?<br>spatially contiguous compartment domains A and B | CDCs comprise a peripheral layer of low compacted, transcriptionally competent CDs and an internal core of more compact, transcriptionally repressed CDs. We propose that chromatin domain clusters (CDCs) correspond with higher order chromatin structures with spatially contiguous A and B compartment domains. This reasoning is consistent with a nanoscale zonation of euchromatic and heterochromatic regions in CDCs. |
| interchromatin compartment (IC) [34,37]<br><br>? | ?<br><br>boundaries [14] | The IC refers to a contiguous 3D channel network, connected to nuclear pore complexes and expanding throughout the nucleus between CDs and CDCs. The finest ramifications of the IC have not yet been defined, but may extend between neighboring nucleosome clutches (NCs) or nanodomains (NDs). Whereas microscopic and Hi-C studies agree with regard to an interchromatin space, comprising ~half of the nuclear space, Hi-C does not allow a quantitative description of the spatial conformation of the IC and its content. Wider IC-lacunas harbor large macromolecular complexes, e.g. splicing speckles. The IC may serve as a ‘road’ system for import of macromolecules, distribution to macromolecular complexes to sites of need, and mRNA export. Boundaries detected in Hi-C experiments represent transition points between contact domains. A role of IC-channels as additional structural boundaries between CDs and CDCs located on both sides, has been considered but not yet proven. |
| perichromatin region (PR) [10,34] | ? | The PR comprises low compacted, transcriptionally competent chromatin lining the IC, and is easily accessible for factors and factor complexes pervading the IC or released from nuclear bodies and splicing speckles. The PR represents the nuclear compartment, where transcription and co-transcriptional splicing of primary transcripts preferentially occurs. Accordingly, the PR is enriched with RNAP II and transcription factories, regulatory and coding sequences of active genes, and epigenetic marks for transcriptionally competent chromatin. |

| | | |
|---|---|---|
| active nuclear compartment (ANC) [34,37]<br><br>inactive nuclear compartment (INC) [34,37] | compartment domains A form chromatin compartment A[15,17,20]<br><br>compartment domains B form chromatin compartment B [15,17,20] | The ANC is formed by the IC together with the PR. We equate transcriptionally competent CDs ('euchromatin') located in the PR and chromatin loops expanding into the IC with compartment domains A, which altogether form the chromatin compartment A. The INC comprises both CDs with repressed genes located in the interior of CDCs ('facultative' heterochromatin) and clusters of 'constitutive' heterochromatin. Like the ANC, the INC is pervaded by IC-channels. Compartment domains A and B are indicated in Hi-C maps by plaid/checkboard patterns. We use the term compartmentalization to describe spatially separated, active and repressed chromatin regions regardless of bin-size. |
| Chromosome territories (CTs) [38] | Enhanced contact frequency within an an entire chromosome [20] | Consistent evidence for a territorial arrangement of chromosomes (CTs) in cell nuclei was obtained in a wide range of animal and plant species both by microscopy and by Hi-C, where contact frequency is often enhanced between all pairs of loci in a single chromosome. |

**References**


1 Natsume, T., Kiyomitsu, T., Saga, Y. & Kanemaki, M. T. Rapid Protein Depletion in Human Cells by Auxin-Inducible Degron Tagging with Short Homology Donors. *Cell Rep* **15**, 210-218, doi:10.1016/j.celrep.2016.03.001 (2016).

2 Wojcik, C. & DeMartino, G. N. Intracellular localization of proteasomes. *Int J Biochem Cell Biol* **35**, 579-589, doi:10.1016/s1357-2725(02)00380-1 (2003).

3 Fei, J. *et al.* Quantitative analysis of multilayer organization of proteins and RNA in nuclear speckles at super resolution. *J Cell Sci* **130**, 4180-4192, doi:10.1242/jcs.206854 (2017).

4 Roukos, V., Pegoraro, G., Voss, T. C. & Misteli, T. Cell cycle staging of individual cells by fluorescence microscopy. *Nat Protoc* **10**, 334-348, doi:10.1038/nprot.2015.016 (2015).

5 Schermelleh, L., Heintzmann, R. & Leonhardt, H. A guide to super-resolution fluorescence microscopy. *J Cell Biol* **190**, 165-175, doi:jcb.201002018 [pii]
10.1083/jcb.201002018 (2010).

6 Miron, E. *et al.* Chromatin arranges in chains of mesoscale domains with nanoscale functional topography independent of cohesin. *bioRxiv566638*, doi:doi.org/10.1101/566638 (2020).

7 Otterstrom, J. *et al.* Super-resolution microscopy reveals how histone tail acetylation affects DNA compaction within nucleosomes in vivo. *Nucleic Acids Res*, doi:10.1093/nar/gkz593 (2019).

8 Ricci, M. A., Manzo, C., Garcia-Parajo, M. F., Lakadamyali, M. & Cosma, M. P. Chromatin fibers are formed by heterogeneous groups of nucleosomes in vivo. *Cell* **160**, 1145-1158, doi:10.1016/j.cell.2015.01.054 (2015).

9 Beagrie, R. A. *et al.* Complex multi-enhancer contacts captured by genome architecture mapping. *Nature* **543**, 519-524, doi:10.1038/nature21411 (2017).

10 Cremer, T. *et al.* Chromosome territories, interchromatin domain compartment, and nuclear matrix: an integrated view of the functional nuclear architecture. *Crit Rev Eukaryot Gene Expr* **10**, 179-212 (2000).

11 Ou, H. D. *et al.* ChromEMT: Visualizing 3D chromatin structure and compaction in interphase and mitotic cells. *Science* **357**, doi:10.1126/science.aag0025 (2017).

12 Zhao, L. *et al.* Chromatin loops associated with active genes and heterochromatin shape rice genome architecture for transcriptional regulation. *Nat Commun* **10**, 3640, doi:10.1038/s41467-019-11535-9 (2019).

13 Zhu, J. J. *et al.* Super resolution imaging of a 1 distinct chromatin loop in human 2 lymphoblastoid cells. *bioRxiv566638*, doi:https://doi.org/10.1101/621920 (2019).

14 Dixon, J. R., Gorkin, D. U. & Ren, B. Chromatin Domains: The Unit of Chromosome Organization. *Mol Cell* **62**, 668-680, doi:10.1016/j.molcel.2016.05.018 (2016).

15 Rao, S. S. *et al.* A 3D map of the human genome at kilobase resolution reveals principles of chromatin looping. *Cell* **159**, 1665-1680, doi:10.1016/j.cell.2014.11.021 (2014).

16 Szabo, Q. *et al.* TADs are 3D structural units of higher-order chromosome organization in Drosophila. *Sci Adv* **4**, eaar8082, doi:10.1126/sciadv.aar8082 (2018).

17 Rowley, M. J. & Corces, V. G. Organizational principles of 3D genome architecture. *Nat Rev Genet* **19**, 789-800, doi:10.1038/s41576-018-0060-8 (2018).

18 Bintu, B. *et al.* Super-resolution chromatin tracing reveals domains and cooperative interactions in single cells. *Science* **362**, doi:10.1126/science.aau1783 (2018).

19 Dixon, J. R. *et al.* Topological domains in mammalian genomes identified by analysis of chromatin interactions. *Nature* **485**, 376-380, doi:10.1038/nature11082 (2012).

20 Lieberman-Aiden, E. *et al.* Comprehensive mapping of long-range interactions reveals folding principles of the human genome. *Science* **326**, 289-293, doi:10.1126/science.1181369 (2009).
21 Fudenberg, G. *et al.* Formation of Chromosomal Domains by Loop Extrusion. *Cell Rep* **15**, 2038-2049, doi:10.1016/j.celrep.2016.04.085 (2016).
22 Sanborn, A. L. *et al.* Chromatin extrusion explains key features of loop and domain formation in wild-type and engineered genomes. *Proc Natl Acad Sci U S A* **112**, E6456-6465, doi:10.1073/pnas.1518552112 (2015).
23 Sexton, T. *et al.* Three-dimensional folding and functional organization principles of the Drosophila genome. *Cell* **148**, 458-472, doi:10.1016/j.cell.2012.01.010 (2012).
24 Baddeley, D. *et al.* Measurement of replication structures at the nanometer scale using super-resolution light microscopy. *Nucleic Acids Res* **38**, e8, doi:10.1093/nar/gkp901 (2010).
25 Ma, H. *et al.* Spatial and temporal dynamics of DNA replication sites in mammalian cells. *J Cell Biol* **143**, 1415-1425 (1998).
26 Marchal, C., Sima, J. & Gilbert, D. M. Control of DNA replication timing in the 3D genome. *Nat Rev Mol Cell Biol*, doi:10.1038/s41580-019-0162-y (2019).
27 Moindrot, B. *et al.* 3D chromatin conformation correlates with replication timing and is conserved in resting cells. *Nucleic Acids Res* **40**, 9470-9481, doi:10.1093/nar/gks736 (2012).
28 Nakamura, H., Morita, T. & Sato, C. Structural organizations of replicon domains during DNA synthetic phase in the mammalian nucleus. *Exp Cell Res* **165**, 291-297, doi:10.1016/0014-4827(86)90583-5 (1986).
29 Pope, B. D. *et al.* Topologically associating domains are stable units of replication-timing regulation. *Nature* **515**, 402-405, doi:10.1038/nature13986 (2014).
30 Sima, J. *et al.* Identifying cis Elements for Spatiotemporal Control of Mammalian DNA Replication. *Cell* **176**, 816-830 e818, doi:10.1016/j.cell.2018.11.036 (2019).
31 Xiang, W. *et al.* Correlative live and super-resolution imaging reveals the dynamic structure of replication domains. *J Cell Biol* **217**, 1973-1984, doi:10.1083/jcb.201709074 (2018).
32 Zhang, H. *et al.* Chromatin structure dynamics during the mitosis-to-G1 phase transition. *Nature* **576**, 158-162, doi:10.1038/s41586-019-1778-y (2019).
33 Zhao, P. A., Rivera-Mulia, J. C. & Gilbert, D. M. Replication Domains: Genome Compartmentalization into Functional Replication Units. *Adv Exp Med Biol* **1042**, 229-257, doi:10.1007/978-981-10-6955-0_11 (2017).
34 Cremer, T. *et al.* The 4D nucleome: Evidence for a dynamic nuclear landscape based on co-aligned active and inactive nuclear compartments. *FEBS Lett* **589**, 2931-2943, doi:10.1016/j.febslet.2015.05.037 (2015).
35 Jackson, D. A. & Pombo, A. Replicon clusters are stable units of chromosome structure: evidence that nuclear organization contributes to the efficient activation and propagation of S phase in human cells. *J Cell Biol* **140**, 1285-1295, doi:10.1083/jcb.140.6.1285 (1998).
36 Cardozo Gizzi, A. M., Cattoni, D. I. & Nollmann, M. TADs or no TADS: Lessons From Single-cell Imaging of Chromosome Architecture. *J Mol Biol* **432**, 682-693, doi:10.1016/j.jmb.2019.12.034 (2020).
37 Cremer, T. *et al.* The Interchromatin Compartment Participates in the Structural and Functional Organization of the Cell Nucleus. *Bioessays* **42**, e1900132, doi:10.1002/bies.201900132 (2020).
38 Cremer, T. & Cremer, M. Chromosome territories. *Cold Spring Harb Perspect Biol* **2**, a003889, doi:10.1101/cshperspect.a003889 (2010).

**Supplementary Table 2: Time lapse imaging data of individual nuclei followed through mitosis**

**Summary:**

| | Control cells | Cohesin depleted cells |
|---|---|---|
| **Total number of mitotic cells** | 45 (100 %) | 36 (100 %) |
| **Inconspicuous mitosis** | 35 (77.8 %) | 4 (11.1 %) |
| **Prolonged mitosis with two daughter cells** | 1 (2.2 %) | - |
| **Prolonged mitosis with formation of MLN** | 5 (11.1 %) | 23 (63.9 %) |
| **Prolonged mitosis not finished within 6 time frames (= 90min)** | 4 (8.9 %) | 9 (25.0 %) |

**Legend:**

**(1) inconspicuous mitosis with two daughter cells (completion within 4 time frames = 60 min).** *Note for cohesin depleted cells: These mitoses were either recorded immediately after addition of auxin (time frame 1) or in cells that did not degrade RAD21-mClover*

**(2) Prolonged mitosis (> 4 time frames) with two daughter cells**

**(3 Prolonged mitosis (> 4 time frames) resulting in one cell with multilobulated nucleus (MLN)**

**(4) Prolonged mitosis not finished within 8 time frames (= 2h):** *Note this fraction includes elongated mitoses that were not finished within the observation period. Since all elongated mitoses in cohesin depleted cells recorded over the entire mitosis resulted in one MLN, the outcome of these mitoses is very likely to be assigned to group 3.*

**Control cells: Single cell data timelapse imaging (time interval 15 minutes)**

| Mitotic cell | Mitosis start (time frame) | Mitosis end (time frame) | Duration [min] | Result |
|---|---|---|---|---|
| 1 | 11 | 14 | 45 | 2 daughter cells |
| 2 | 12 | 16 | 60 | 2 daughter cells |
| 3 | 46 | 50 | 60 | 2 daughter cells |
| 4 | 50 | 54 | 60 | 2 daughter cells |
| 5 | 21 | 24 | 45 | 2 daughter cells |
| 6 | 21 | 25 | 60 | 2 daughter cells |
| 7 | 23 | 79 | 840 | 1 cell with MLN |
| 8 | 33 | 62 | 435 | 1 cell with MLN |
| 9 | 10 | 13 | 45 | 2 daughter cells |
| 10 | 11 | 14 | 45 | 2 daughter cells |
| 11 | 16 | 20 | 60 | 2 daughter cells |
| 12 | 19 | 24 | 75 | 2 daughter cells |
| 13 | 20 | 23 | 45 | 2 daughter cells |
| 14 | 39 | 68 | 435 | 1 cell with MLN |
| 15 | 70 | >84 | >210 | Mitosis not finished |
| 16 | 73 | 77 | 60 | 2 daughter cells |
| 17 | 78 | 82 | 60 | 2 daughter cells |
| 18 | 1 | 4 | 60 | 2 daughter cells |
| 19 | 12 | 17 | 60 | 2 daughter cells |
| 20 | 23 | 27 | 60 | 2 daughter cells |
| 21 | 25 | 28 | 45 | 2 daughter cells |
| 22 | 4 | 8 | 60 | 2 daughter cells |
| 23 | 8 | 11 | 45 | 2 daughter cells |
| 24 | 9 | 12 | 45 | 2 daughter cells |
| 25 | 11 | 14 | 45 | 2 daughter cells |
| 26 | 74 | >84 | >150 | Mitosis not finished |
| 27 | 32 | 36 | 60 | 2 daughter cells |
| 28 | 45 | 49 | 60 | 2 daughter cells |
| 29 | 53 | 78 | 375 | 1 cell with MLN |
| 30 | 3 | 7 | 60 | 2 daughter cells |
| 31 | 25 | 29 | 60 | 2 daughter cells |
| 32 | 31 | 34 | 45 | 2 daughter cells |
| 33 | 39 | 42 | 45 | 2 daughter cells |
| 34 | 44 | 47 | 45 | 2 daughter cells |
| 35 | 46 | 49 | 45 | 2 daughter cells |

| | | | | |
|---|---|---|---|---|
| 36 | 8 | 12 | 60 | 2 daughter cells |
| 37 | 14 | 18 | 60 | 2 daughter cells |
| 38 | 13 | 34 | 315 | 1 cell with MLN |
| 39 | 16 | 32 | 240 | 2 daughter cells |
| 40 | 65 | 70 | 75 | 2 daughter cells |
| 41 | 70 | >84 | >210 | Mitosis not finished |
| 42 | 3 | 7 | 60 | 2 daughter cells |
| 43 | 5 | 8 | 45 | 2 daughter cells |
| 44 | 11 | 14 | 45 | 2 daughter cells |
| 45 | 60 | >84 | >360 | Mitosis not finished |

**Cohesin depleted cells: Single cell data timelapse imaging (time interval 15 minutes)**

| Mitotic cell | Mitosis start (time frame) | Mitosis end (time frame) | Duration [min] | Result | RAD21-mClover Fluorescence ++ + - |
|---|---|---|---|---|---|
| 1 | 1 | 5 | 75 | 2 daughter cells | ++ |
| 2 | 9 | 29 | 300 | 1 cell with MLN | - |
| 3 | 12 | >44 | >480 | Mitosis not finished | |
| 4 | 59 | 77 | 270 | 1 cell with MLN | - |
| 5 | 58 | 82 | 360 | 1 cell with MLN | - |
| 6 | 59 | 84 | 375 | 1 cell with MLN | - |
| 7 | 1 | 5 | 75 | 2 daughter cells | + |
| 8 | 13 | 30 | 255 | 1 cell with MLN | - |
| 9 | 14 | 53 | 585 | 1 cell with MLN | - |
| 10 | 36 | 50 | 210 | 1 cell with MLN | - |
| 11 | 41 | 53 | 180 | 1 cell with MLN | - |
| 12 | 14 | 51 | 555 | 1 cell with MLN | - |
| 13 | 18 | >60 | >630 | Mitosis not finished | - |
| 14 | 30 | >61 | >465 | Mitosis not finished | - |
| 15 | 36 | >64 | >420 | Mitosis not finished | - |
| 16 | 44 | 62 | 270 | 1 cell with MLN | - |
| 17 | 48 | 63 | 225 | 1 cell with MLN | - |
| 18 | 66 | >84 | >270 | Mitosis not finished | - |
| 19 | 69 | >84 | >225 | Mitosis not finished | - |
| 20 | 73 | >84 | >165 | Mitosis not finished | - |
| 21 | 74 | >84 | >150 | Mitosis not finished | - |
| 22 | 13 | >58 | >675 | Mitosis not finished | - |
| 23 | 50 | 72 | 330 | 1 cell with MLN | - |
| 24 | 55 | 73 | 270 | 1 cell with MLN | - |
| 25 | 13 | 50 | 555 | 1 cell with MLN | - |
| 26 | 27 | 82 | 825 | 1 cell with MLN | - |
| 27 | 28 | 44 | 240 | 1 cell with MLN | - |
| 28 | 39 | 51 | 180 | 1 cell with MLN | - |
| 29 | 40 | 62 | 330 | 1 cell with MLN | - |
| 30 | 52 | 66 | 210 | 1 cell with MLN | - |
| 31 | 40 | 62 | 330 | 1 cell with MLN | - |
| 32 | 47 | 61 | 210 | 1 cell with MLN | - |
| 33 | 60 | 83 | 345 | 1 cell with MLN | - |
| 34 | 13 | 17 | 60 | 2 daughter cells | ++ (escaper) |
| 35 | 17 | 21 | 60 | 2 daughter cells | ++(escaper) |
| 36 | 42 | 59 | 255 | 1 cell with MLN | - |

**Supplementary Table 3: Significance tests related to Figs. 3, 4 and 8**
p-values ($p < 0.05$ highlighted in light gray, < 0.01 in dark gray) of data shown in Fig. 3D (DAPI intensity classes) are listed in 'DAPI' columns. p-values of data shown in Fig. 4G-H (distribution of SC35, H3K27me3 and RNA Pol II signals on DAPI intensity classes) are listed in the respective rows. Test: Mann-Whitney test (Wilcoxon rank sum test). Software: RStudio (Version 1.0.143). Correction: Bonferroni-Holm correction for multiple tests

| Group | n = cell number | | | |
|---|---|---|---|---|
| | DAPI | Pol II | SC35 | H3K27me3 |
| control | 38 | 20 | 18 | 18 |
| 6h auxin | 39 | 17 | 22 | 22 |
| 30h auxin | 33 | 16 | 17 | 17 |

**Class 1**

| Tested groups | p-value | | | |
|---|---|---|---|---|
| | DAPI | Pol II | SC35 | H3K27me3 |
| control ↔ 6h auxin | 0.884 | 1 | 0.413 | 1 |
| control ↔ 30h auxin | 0.788 | 1 | 0.414 | 0.002 |
| 6h auxin ↔ 30h auxin | 0.012 | 1 | 0.117 | 0.043 |

**Class 2**

| Tested groups | p-value | | | |
|---|---|---|---|---|
| | DAPI | Pol II | SC35 | H3K27me3 |
| control ↔ 6h auxin | 0.950 | 1 | 0.384 | 0.435 |
| control ↔ 30h auxin | 0.680 | 1 | 0.613 | 0.032 |
| 6h auxin ↔ 30h auxin | 0.680 | 1 | 0.064 | 0.119 |

**Class 3**

| Tested groups | p-value | | | |
|---|---|---|---|---|
| | DAPI | Pol II | SC35 | H3K27me3 |
| control ↔ 6h auxin | 0.242 | 1 | 0.206 | 1 |
| control ↔ 30h auxin | 0.963 | 1 | 0.618 | 1 |
| 6h auxin ↔ 30h auxin | 0.014 | 1 | 0.118 | 1 |

**Class 4**

| Tested groups | p-value | | | |
|---|---|---|---|---|
| | DAPI | Pol II | SC35 | H3K27me3 |
| control ↔ 6h auxin | 1 | 1 | 0.399 | 1 |
| control ↔ 30h auxin | 0.801 | 1 | 0.532 | 0.46 |
| 6h auxin ↔ 30h auxin | 0.070 | 1 | 0.136 | 1 |

**Class 5**

| Tested groups | p-value | | | |
|---|---|---|---|---|
| | DAPI | Pol II | SC35 | H3K27me3 |
| control ↔ 6h auxin | 1 | 1 | 0.470 | 0.794 |
| control ↔ 30h auxin | 1 | 1 | 0.455 | 0.028 |
| 6h auxin ↔ 30h auxin | 0.360 | 1 | 0.135 | 0.042 |

**Class 6**

| Tested groups | p-value | | | |
|---|---|---|---|---|
| | DAPI | Pol II | SC35 | H3K27me3 |
| control ↔ 6h auxin | 1 | 1 | 0.398 | 0.823 |
| control ↔ 30h auxin | 0.041 | 1 | 0.085 | 0.031 |
| 6h auxin ↔ 30h auxin | 0.132 | 1 | 0.010 | 0.036 |

**Class 7**

| Tested groups | p-value | | | |
|---|---|---|---|---|
| | DAPI | Pol II | SC35 | H3K27me3 |
| control ↔ 6h auxin | 1 | 1 | 0.487 | 1 |
| control ↔ 30h auxin | 0.800 | 1 | 0.006 | 0.030 |
| 6h auxin ↔ 30h auxin | 0.480 | 1 | 0.0001 | 0.050 |

**Standard deviation for DAPI signal distribution in classes**

| Class | 1 | 2 | 3 | 4 | 5 | 6 | 7 |
|---|---|---|---|---|---|---|---|
| **control** | 0.0561 | 0.0160 | 0.0226 | 0.0227 | 0.0173 | 0.0086 | 0.0028 |
| **6h auxin** | 0.0423 | 0.0808 | 0.0321 | 0.0236 | 0.0171 | 0.0105 | 0.0045 |
| **30h auxin** | 0.0327 | 0.0260 | 0.0149 | 0.0228 | 0.0187 | 0.0096 | 0.0036 |

**Standard deviation for SC35 signal distribution in classes**

| Class | 1 | 2 | 3 | 4 | 5 | 6 | 7 |
|---|---|---|---|---|---|---|---|
| **control** | 0.1302 | 0.1034 | 0.0301 | 0.0101 | 0.0030 | 0.0008 | 0.0001 |
| **6h auxin** | 0.0862 | 0.0559 | 0.0240 | 0.0082 | 0.0023 | 0.0005 | 0.000 |
| **30h auxin** | 0.1674 | 0.0846 | 0.0509 | 0.0240 | 0.0103 | 0.0037 | 0.0014 |

**Standard deviation for H3K27me3 signal distribution in classes**

| Class | 1 | 2 | 3 | 4 | 5 | 6 | 7 |
|---|---|---|---|---|---|---|---|
| **control** | 0.0438 | 0.0453 | 0.0434 | 0.0385 | 0.0500 | 0.0274 | 0.0034 |
| **6h auxin** | 0.0465 | 0.0509 | 0.0473 | 0.0399 | 0.0688 | 0.0241 | 0.0042 |
| **30h auxin** | 0.0294 | 0.0699 | 0.0452 | 0.0433 | 0.0457 | 0.0588 | 0.0112 |

**Standard deviation for Pol II Ser5P signal distribution in classes**

| Class | 1 | 2 | 3 | 4 | 5 | 6 | 7 |
|---|---|---|---|---|---|---|---|
| **control** | 0.1226 | 0.0687 | 0.0622 | 0.0747 | 0.0383 | 0.0179 | 0.0021 |
| **6h auxin** | 0.1506 | 0.1052 | 0.0738 | 0.0894 | 0.0518 | 0.0173 | 0.0021 |
| **30h auxin** | 0.1228 | 0.1014 | 0.0878 | 0.0884 | 0.0337 | 0.0092 | 0.0022 |

### Average nuclear volume: significance between series (Fig. 3)

Test: Mann-Whitney test of data shown in Fig. 3E
Software: Addinsoft (2019). XLSTAT statistical and data analysis solution. Boston, USA. https://www.xlstat.com. (Version 2019.2.3, Build 59941); Significant: $p < 0.05$

| Tested group | p-value |
|---|---|
| control ↔ 6h auxin | 0.402 |
| control ↔ 30h auxin | <0.0001 |
| 6h auxin ↔ 30h auxin | <0.0001 |

### RD volume: significance between series (Fig. 8)

Test: Mann-Whitney test of data shown in Fig. 8C
Software: RStudio (Version 1.0.143).
Correction: Bonferroni-Holm correction for multiple tests; Significant: $p < 0.05$

| Tested group | p-value |
|---|---|
| control ↔ 6h auxin | <0.0001 |
| control ↔ 30h auxin | <0.0001 |
| 6h auxin ↔ 30h auxin | <0.0001 |
